# *SNAP18* Truncation Triggers a Competitive Binding Switch Between NSF and ATG8f, Balancing Vesicular Trafficking and Autophagy for SCN Resistance in Soybean

**DOI:** 10.64898/2026.02.24.707681

**Authors:** Dongmei Wang, Liang Wang, Qiannan Liu, Luying Chen, Lin Weng, Hui Yu, Chunjie Li, Minghui Huang, Suxin Yang, Xianzhong Feng, Shaojie Han

## Abstract

Soybean cyst nematode (SCN, *Heterodera glycines*) poses the most devastating biotic threat to global soybean production. Traditional SCN resistance mediated by the *Rhg1* locus predominantly relies on gene copy number expansion and elevated α-SNAP protein dosage. Here, we report a novel resistance mechanism in a single-copy *rhg1-c* background, wherein a C-terminal 24-amino-acid truncation of SNAP18 (designated SNAP18_lmm3_) triggers a functional switch from vesicular trafficking to autophagic degradation. Biochemical assays and structural modeling demonstrate that this truncation severely impairs the canonical interaction between SNAP18 and N-ethylmaleimide-sensitive factor (NSF), disrupting SNARE complex recycling and inducing localized cytotoxicity. Concomitantly, the truncated SNAP18_lmm3_ exposes a binding interface for the autophagy-related protein ATG8f, routing the aberrant protein for selective autophagic clearance. This constitutively activated autophagic flux acts as a systemic detoxification system, preventing widespread cell death and ensuring normal plant growth under non-stressed conditions. Upon SCN infection, SNAP18_lmm3_ specifically hyper-accumulates within nematode-induced syncytia. This accumulation reaches levels fourfold higher than in adjacent cells, which overwhelms the local autophagic capacity and triggering targeted cell death that arrests nematode development. By elucidating this competitive molecular switch between NSF and ATG8f binding, our study establishes a “self-degrading toxin” model that resolves the inherent trade-off between plant growth and immunity. This work provides a new theoretical framework for engineering cellular homeostasis to enhance durable crop resistance against parasitic nematodes.

## Introduction

Soybean cyst nematode (SCN, *Heterodera glycines* Ichinohe) remains the most devastating biotic threat to global soybean production, causing yield losses that exceed US$1.5 billion annually in the United States alone (Bandara et al. 2020; Bent 2022; Jones et al. 2013). The sedentary endoparasitic lifestyle of SCN involves a sophisticated manipulation of host cellular processes to induce specialized, multinucleated feeding structures called syncytia within the root vascular cylinder (Yusuf and Bello 2025; Yusuf et al. 2026). These syncytia serve as the sole metabolic sink for the developing nematode, requiring massive reprogramming of host vesicular trafficking and nutrient flow (Bent 2022). For decades, soybean breeding has relied heavily on the ’Resistance to *Heterodera glycines* 1’ (*Rhg1*) locus to manage this pathogen (Cook et al. 2014; Cook et al. 2012). However, the continuous and intensive use of limited genetic resources has led to the widespread emergence of virulent SCN populations, necessitating the discovery of novel genetic determinants and a deeper mechanistic understanding of the cell biology underlying SCN-soybean interactions (Chen et al. 2021; Meinhardt et al. 2021; Niblack et al. 2008; Peng et al. 2021).

The *Rhg1* locus consists of a tandemly repeated genomic block containing four genes, among which *SNAP18* (encoding an α-soluble NSF attachment protein, *α-SNAP*) is the primary determinant of resistance (Cook et al. 2012; Dong and Hudson 2022; Han et al. 2023; Lakhssassi et al. 2017; Lee et al. 2015). *Rhg1*-mediated immunity is inherently dosage-dependent; copy-number expansion significantly increases the transcript levels of atypical *α-SNAP* isoforms in resistant accessions (Cook et al. 2014; Cook et al. 2012; Patil et al. 2019). Well-characterized variants include the *rhg1-b* haplotype from PI 88788 (high-copy) and the *rhg1-a* haplotype from Peking (low-copy), the latter requiring epistatic interaction with the *Rhg4* locus for full resistance (Bayless et al. 2019; Liu et al. 2012). In contrast, the vast majority of soybean cultivars carry only a single copy of the canonical *SNAP18* (the *rhg1-c* haplotype) and remain highly susceptible to SCN infection (Cook et al. 2014). Despite the central importance of SNAP18s in nematode defense, identifying potent resistance alleles in single-copy backgrounds has remained an elusive goal for soybean improvement (Suzuki 2020; Usovsky et al. 2023).

Canonical SNAP18s physically interact with the vesicle-trafficking component N-ethylmaleimide-sensitive factor (NSF), facilitating the disassembly and reuse of SNARE (soluble NSF-attachment protein receptor) complexes during vesicle docking and fusion (Bombardier and Munson 2015). Variation in α-SNAPs can impact their interactions with NSF and disrupt the SNARE complex; atypical SNAP18s impede syncytium and nutrient flow, leading to SCN resistance (Bayless et al. 2016). SCN-resistant soybeans possess normal SNAP18s in other contexts, which can offset the effects of *Rhg1* variants. Certain NSF variants, such as NSF_RAN07_, have improved functionality when interacting with *Rhg1*-encoded atypical *α-SNAPs*, reducing cytotoxicity and promoting soybean growth (Bayless et al. 2018). Atypical *α-SNAPs* are found in multi-copy *Rhg1* germplasms, but their specific function in nematode resistance remains unclear, and none have been identified in single-copy *Rhg1* varieties to date.

Autophagy is a vital process that degrades cellular components within lysosomes and vacuoles, playing a role in cellular stability, survival during nutrient scarcity, and responses to harmful stimuli. Over 40 *AUTOPHAGY-RELATED* (*ATG*) genes have been identified in plants, each contributing to various stages of autophagy, including induction, phagosome formation, extension, autophagosome maturation and vacuole fusion (Marshall and Vierstra 2018; Tang and Bassham 2018). Despite these advancements, the understanding of autophagy’s role in resistance to plant-parasitic nematodes remains limited. Cyst nematodes use the effector protein NMAS1 to interact with plant autophagy machinery, facilitating their infection (Chen et al. 2023). Autophagy helps tomato defend against root-knot nematodes by enhancing jasmonate-related immunity (Zou et al. 2023). However, whether autophagy serves as a regulatory bridge that balances the cytotoxicity of vesicle trafficking components with nematode resistance has not been investigated in soybean.

In this study, we identified the *lesion-mimic mutant 3* (*lmm3*) in the SCN-susceptible Williams 82 background, which encodes a truncated SNAP18 isoform (SNAP18_lmm3_) with a 24-amino-acid C-terminal deletion. We demonstrate that this truncation disrupts the canonical SNAP18-NSF interaction and triggers a constitutive activation of autophagic flux to mitigate the resulting cytotoxicity. Our findings reveal that SNAP18_lmm3_ acts as a “self-degrading toxin” that hyper-accumulates specifically at SCN feeding sites to arrest nematode development while being systemically cleared by ATG8f-mediated autophagy to ensure plant survival. This discovery unravels a sophisticated biochemical switch of SNAP18 between vesicular trafficking and autophagic degradation, providing a novel framework for engineering durable crop immunity through cellular homeostasis.

## Results

### C-Terminal Truncation of SNAP18_lmm3_ Disrupts its Canonical Interaction with NSF

To investigate the molecular mechanisms of soybean resistance to SCN in single-copy *Rhg1* environments, we identified a lesion-mimic mutant, designated as *lmm3*, from an ethyl methanesulfonate (EMS)-mutagenized population of the SCN-susceptible cultivar Williams 82 (*rhg1-c*). The *lmm3* mutant exhibits spontaneous rust-brown necrotic lesions on its leaves and demonstrates significantly enhanced resistance to SCN infection. Through map-based cloning, we determined that the *lmm3* phenotype is caused by a G-to-A transition at nucleotide 797 of the *α-SNAP* gene *SNAP18* (*Glyma.18G022500)*. This mutation introduces a premature stop codon (Trp266*), resulting in a 24-amino-acid truncation at the C-terminus of the encoded protein (hereafter referred to as SNAP18_lmm3_) (Figure 1A). The evolutionarily conserved C-terminal helical bundle of *α-SNAP* is essential for its physical association with the N-terminal domain of NSF. In atypical *α-SNAPs*, specifically SNAP_LC_ (encoded by the low-copy *rhg1-a* locus) and SNAP_HC_ (encoded by the high-copy *rhg1-b* locus), the disruption of this domain leads to attenuated NSF binding (Lakhssassi et al. 2017; Matsye et al. 2012). This failure to assemble a functional complex impairs vesicular trafficking and triggers localized cytotoxicity, providing the mechanistic basis for the arrest of nematode development (Bayless et al. 2016). Since SNAP18_lmm3_ lacks a critical portion of this interaction interface, we hypothesized that its ability to form a functional 20S complex with NSF would be severely compromised.

**FIGURE 1.**
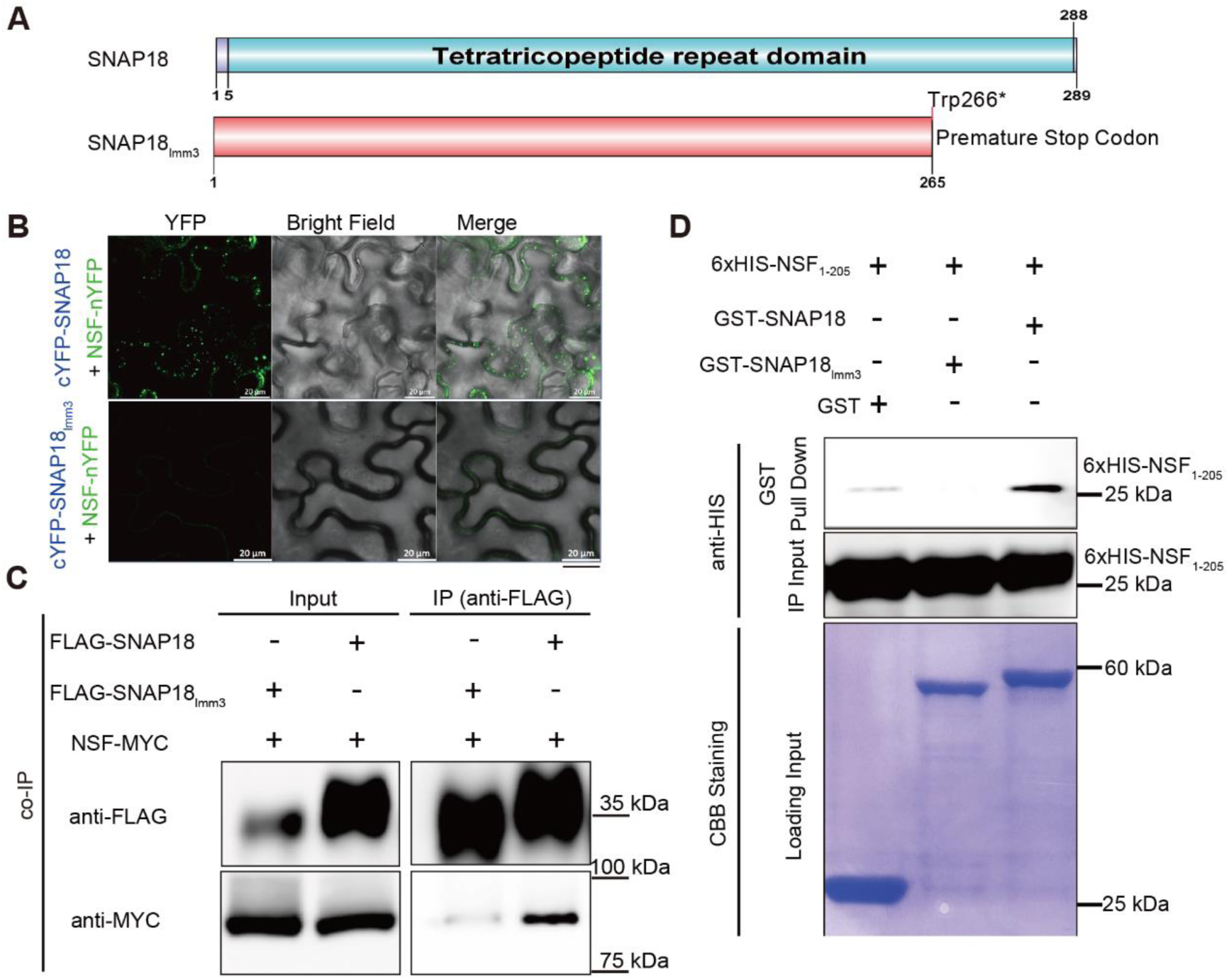
C-terminal truncation of SNAP18lmm3 disrupts its canonical interaction with NSF. (A) Schematic representation of the domain architecture of wild-type SNAP18 and the truncated mutant SNAP18lmm3. (B) BiFC assay demonstrating weakened binding of SNAP18lmm3 to NSF. Representative images show transient co-expression of NSF-nYFP with N-terminal cYFP-tagged SNAP18 and SNAP18lmm3 in *N. benthamiana* leaves. Scale bars = 20 μm. (C) Co-immunoprecipitation of SNAP18lmm3 with NSF in *N. benthamiana* leaves. NSF*–*MYC was detected at lower levels in the immunoprecipitants of FLAG*–*SNAP18lmm3 compared to FLAG*–*SNAP18. (D) Pull-down of 6 × His*–*NSF1-205 with GST-tagged SNAP18 and SNAP18lmm3. Only GST*–*SNAP18 was observed to bind with 6 ×His*–*NSF1-205, while no interaction between GST*–*SNAP18lmm3 and 6 ×His*–*NSF was detected.

To test this, we performed bimolecular fluorescence-complementation (BiFC) and co-immunoprecipitation (co-IP) experiments to investigate potential interactions between SNAP18_lmm3_ and NSF. Fluorescent vesicles appeared at the plasma membrane when NSF-nYFP and cYFP-SNAP18 were co-expressed, indicating that the interaction between SNAP18 and NSF plays a role in vesicle formation at the plasma membrane (Figure 1B; Figure S1). We observed weakened interactions between SNAP18_HC_ or SNAP18_LC_ with NSF, due to C-terminal amino-acid changes (Matsye et al. 2012), as evidenced by fewer fluorescent vesicles visible on the plasma membrane (Figure S1). The complete disappearance of fluorescent vesicles was observed upon co-expression of SNAP18_lmm3_ and NSF (Figure 1B; Figure S1), suggesting an alteration in the interaction between SNAP18_lmm3_ and NSF. Co-IP analysis indicated a substantial decrease in NSF binding to SNAP18_lmm3_ compared to SNAP18 (Figure 1C). Furthermore, the C-terminal-truncated SNAP18 (SNAP18_lmm3_) exhibited impaired interaction with the N-terminus of NSF in a pull-down assay (Figure 1D). These findings highlight compromised binding capacity between SNAP18_lmm3_ and NSF (Figure 1 B-D).

### Impaired NSF-SNAP18 Interaction Triggers Vesicular Trafficking Arrest and Cytotoxicity

SNAP18 proteins are known to influence plant-cell death through vesicle trafficking and membrane fusion (Matsye et al. 2012). Cell-death-induction assays in *N. benthamiana* demonstrated cytotoxicity for SNAP18_lmm3_, though this induction was slower and less severe than SNAP18_HC/LC_ (Figure 2A). This effect was not observed with wild-type α-SNAP/SNAP18. Furthermore, to evaluate whether these variants disrupted intracellular trafficking, we used a GFP-secretion assay (Bayless et al. 2016). This system employs an engineered secreted GFP (secGFP), which fluoresces strongly when retained in the ER-Golgi network due to trafficking disruptions, but fluoresces weakly when successfully secreted to the apoplast. Co-expression of SNAP18_lmm3_ strongly disrupted secGFP trafficking, as evidenced by intense intracellular GFP fluorescence, while SNAP18_HC_ and SNAP18_LC_ caused partial disruptions (Figure 2B). In contrast, wild-type SNAP18 did not affect trafficking, showing very weak fluorescence. These results suggest that the failure to recycle SNARE complexes via NSF leads to a “trafficking paralysis” that underpins the *lmm3* autoimmune phenotype.

**FIGURE 2.**
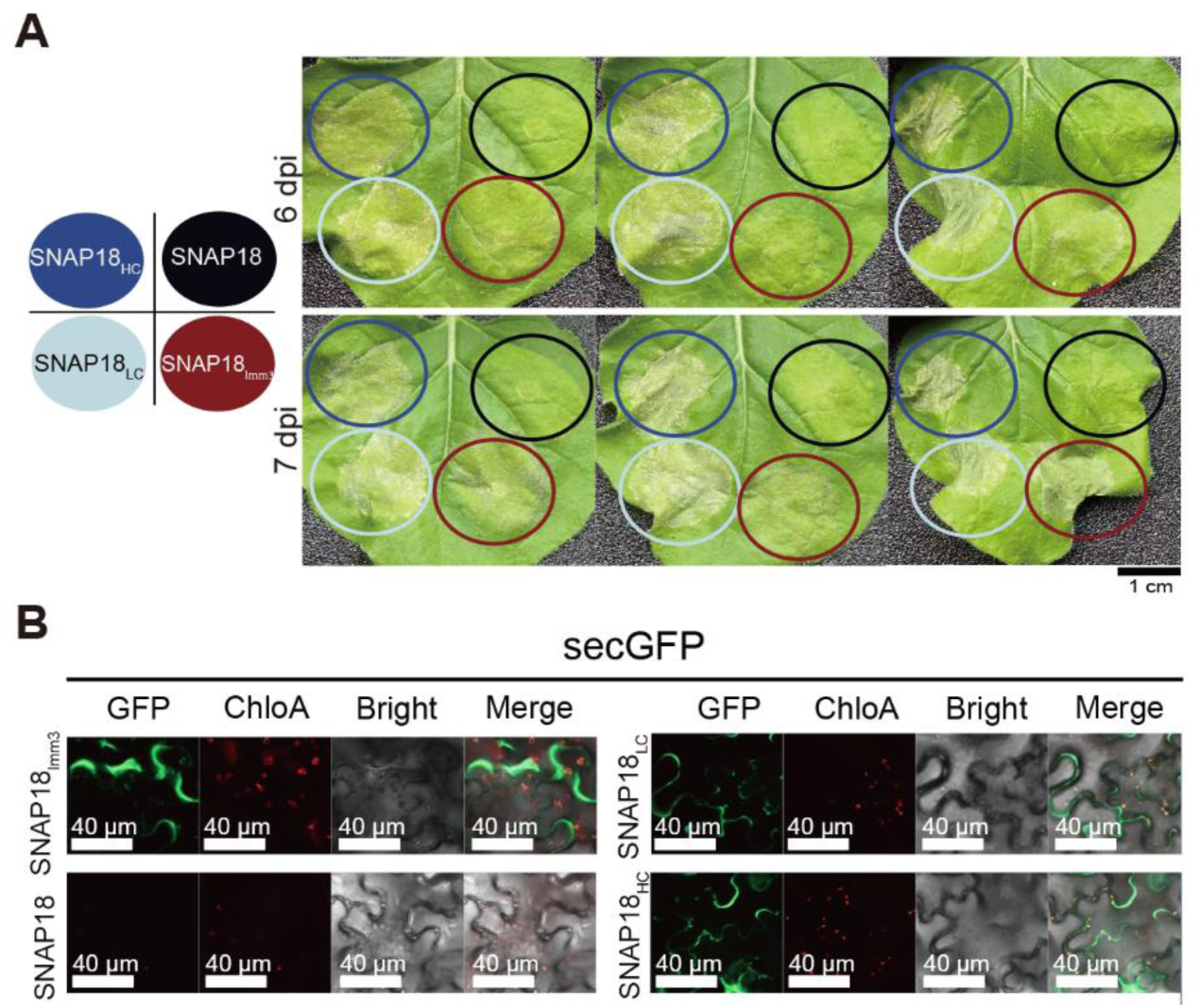
Impaired NSF-SNAP18 interaction triggers vesicular trafficking arrest and cytotoxicity. (A) Cell-death assays in *N. benthamiana* leaves. Left: Schematic of leaf zones infiltrated with the indicated constructs. Right: Images of leaves transiently expressing untagged SNAP18 isoforms at 6 and 7 days after infiltration. Three biological replicates are shown. Scale bar = 1 cm. (B) Confocal-microscopy analysis of secGFP assay. Green fluorescence indicates secGFP retention in the ER–Golgi, while weak fluorescence indicates secretion to the apoplast. SNAP18lmm3, SNAP18HC and SNAP18LC caused strong intracellular retention and wild-type SNAP18 showed no obvious disruption. Red fluorescence marks chloroplast autofluorescence (ChloA). Panels: GFP, ChloA, bright-field, and merged views. Scale bars: 40 μm.

### SCN Infection Induces Localized Hyper-accumulation of SNAP18_lmm3_

Next, we examined whether SNAP18_lmm3_ was differentially expressed upon SCN challenge. Roots of Williams 82, *lmm3*, and two SCN-resistant cultivars (PI88788 and Forrest) were inoculated and then assayed by western blot using a custom α-SNAP antibody. Increases in SNAP18_lmm3_ abundance were seen in SCN-infected areas at 3 dpi, indicating induction upon infection (Figure 3A,B). *SNAP18* transcript abundance remained stable at 1 dpi and 3 dpi in both Williams 82 and *lmm3* (Figure S2A,B). Immunogold labeling of SNAP18_lmm3_ and electron microscopy indicated accumulation primarily in syncytia at 4 dpi, four times greater than adjacent cells (Figure 3C,D and Figure S2C). This pattern of increased syncytial localization aligns with hyper accumulation of SNAP18 variants SNAP18_LC_ and SNAP18_HC_. In summary, SNAP18_lmm3_ specifically accumulates within syncytia upon SCN infection, similar to SNAP18_HC_ and SNAP18_LC_.

**FIGURE 3.**
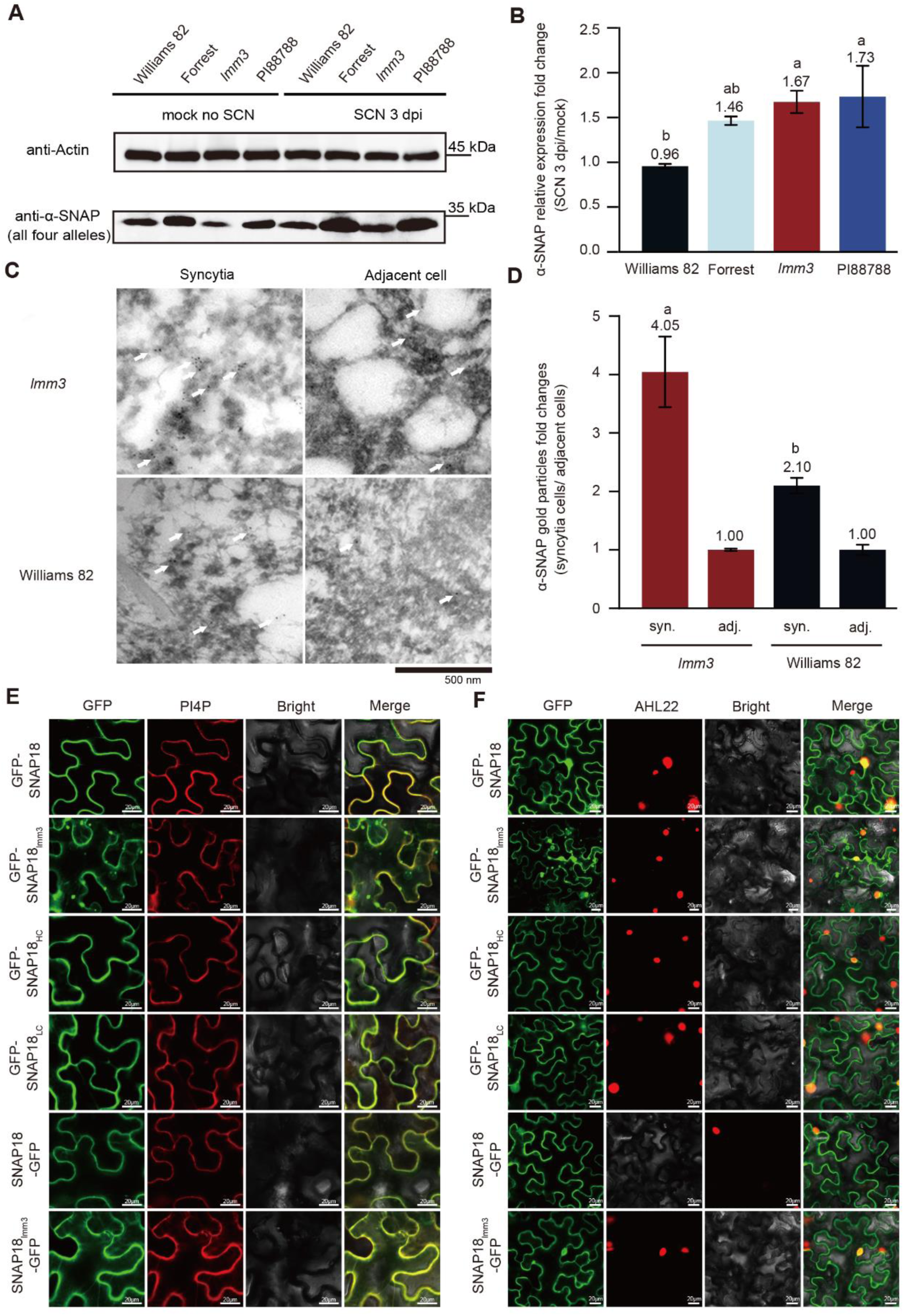
(A) Western blot of α-SNAP abundance in root samples from SCN-infected and un-infected Williams 82, Forrest, *lmm3* and PI88788 harvested 3 dpi. Actin was used as a loading control. (B) Relative α-SNAP expression based on gray-value analysis of bands from western blotting on (A). Data are the mean ±SEM of n = 3. One-way ANOVA, different letters represent *P* < 0.05. (C) Representative electron micrographs displaying the accumulation of immunogold-labeled SNAP18 and SNAP18lmm3 in syncytial cells at 4 dpi. Immunogold-labeled α-SNAPs (visible as solid black particles and indicated by white arrows were imaged in syncytia cells and in adjacent cells from root samples. (D) Quantification of α-SNAPs immunogold particles in syncytial (syn.) cells, in comparison to adjacent (adj.) cells on the same grid in (C). This analysis was based on at least 18 images from three independent experiments. Data are the mean ±SEM of n ≥ 2, two-way ANOVA, *P* < 0.05, different letters represent statistically significant differences. (E) Transient expression in *N. benthamiana* leaves of N-terminally GFP-tagged SNAP18 isoforms SNAP18, SNAP18HC, SNAP18LC and SNAP18lmm3 (upper four panels) and C-terminally GFP-tagged SNAP18 and SNAP18lmm3 (lower two panels). *At*PI4P–mCherry was used to mark plasma-membrane localization. Scale bar = 20 μm. (F) Transient expression in *N. benthamiana* leaves of N-terminally GFP-tagged SNAP18 isoforms, including SNAP18, SNAP18HC, SNAP18LC and SNAP18lmm3 (upper four panels) and C-terminally GFP-tagged SNAP18 and SNAP18lmm3 (lower two panels). *At*AHL22–mCherry was to mark nuclear localization. Scale bar = 20 μm.

To better understand the function of SNAP18_lmm3_, we tracked subcellular localization by fusing GFP to either the N-terminus or C-terminus and transiently expressing the fusion proteins in *N. benthamiana* leaves. Interestingly, in addition to its presence on the plasma membrane (similar to SNAP18, SNAP18_HC_ and SNAP18_LC_), GFP-SNAP18_lmm3_ fusions displayed a distinctive vesicle-like localization pattern (Figure 3E,F). However, SNAP18_lmm3_-GFP did not replicate this pattern (Figure 3E,F). The exposure of the C-terminus appears crucial for the vesicular localization of SNAP18_lmm3_, indicating its importance in determining proper subcellular localization and thus function (Figure 3E,F). This vesicle-like pattern of GFP-SNAP18_lmm3_ may suggest vesicle retention within the *N. bethamiana* cells, disrupting normal vesicle trafficking.

In summary, SNAP18_lmm3_, akin to other aberrant SNAP18s (SNAP18_HC/LC_), compromises proper NSF function, is likely to impair proper NSF function, potentially disrupting vesicle trafficking and contributing to cytotoxicity. It is up-regulated in SCN-infected root regions and accumulates in syncytia at SCN feeding sites. Notably, *SNAP18_lmm3_* is a mutant allele in the Williams 82 background, a susceptible genotype that lacks the SCN resistance loci *Rhg1* and *Rhg4*.

### Autophagy Mediates the Systemic Detoxification of SNAP18_lmm3_

Despite the inherent cytotoxicity of SNAP18_lmm3_ and the resulting spontaneous necrosis, the *lmm3* mutant remains viable and capable of producing seeds (Wang et al. accompanying manuscript). This survival is remarkable given that in typical *Rhg1* lines (e.g., PI 88788), the detoxification of aberrant SNAP18 variants (SNAP_HC/LC_) necessitates a specialized NSF isoform, NSF_RAN07_, to mitigate protein-induced toxicity (Bayless et al. 2018). Since *lmm3* was identified in the single-copy *rhg1-c* background of Williams 82, which lacks such compensatory NSF variants, we hypothesized that an alternative cellular pathway must exist to manage the accumulation of the truncated SNAP18_lmm3_ protein.

To test this, we first examined steady-state SNAP18 abundance using a custom anti-SNAP18 antibody that recognizes the conserved N-termini of all SNAP18 alleles. Western blot analysis of leaves and roots at 7 days after germination (DAG) revealed that total SNAP18 protein levels were significantly lower in *lmm3* compared to Williams 82, Forrest, and PI 88788 (Figure 4A,B), while transcript levels were statistically non-significantly different between Williams 82 and *lmm3* roots (Figure S2A,B). This suggests that differences in protein abundance likely arise from post-transcriptional regulation. To investigate this further, we transiently expressed *SNAP18*, *SNAP18_lmm3_*, *SNAP18_HC_*, and *SNAP18_LC_* under the constitutive *35S* promoter in *N. benthamiana*. We observed less SNAP18_lmm3_ compared to other isoforms (Figure S3), consistent with findings in soybean (Figure 4A,B). We hypothesized that SNAP18_lmm3_ degradation might involve the 26S ubiquitin-proteasome pathway and therefore treated infiltrated leaves with the proteasome inhibitor MG132. No obvious change was seen in SNAP18_lmm3_ abundance compared to controls, suggesting that its degradation is not mediated by the 26S proteasome (Figure S3). Next, we transiently expressed FLAG-tagged SNAP18 isoforms followed by treatment with MG132 or the autophagy inhibitor E-64-D. Anti-FLAG western blots indicated that E-64-D treatment could stabilize SNAP18_lmm3_, while MG132 treatment again had no obvious effect (Figure 4C). Thus, SNAP18_lmm3_ may undergo degradation via autophagy.

**FIGURE 4.**
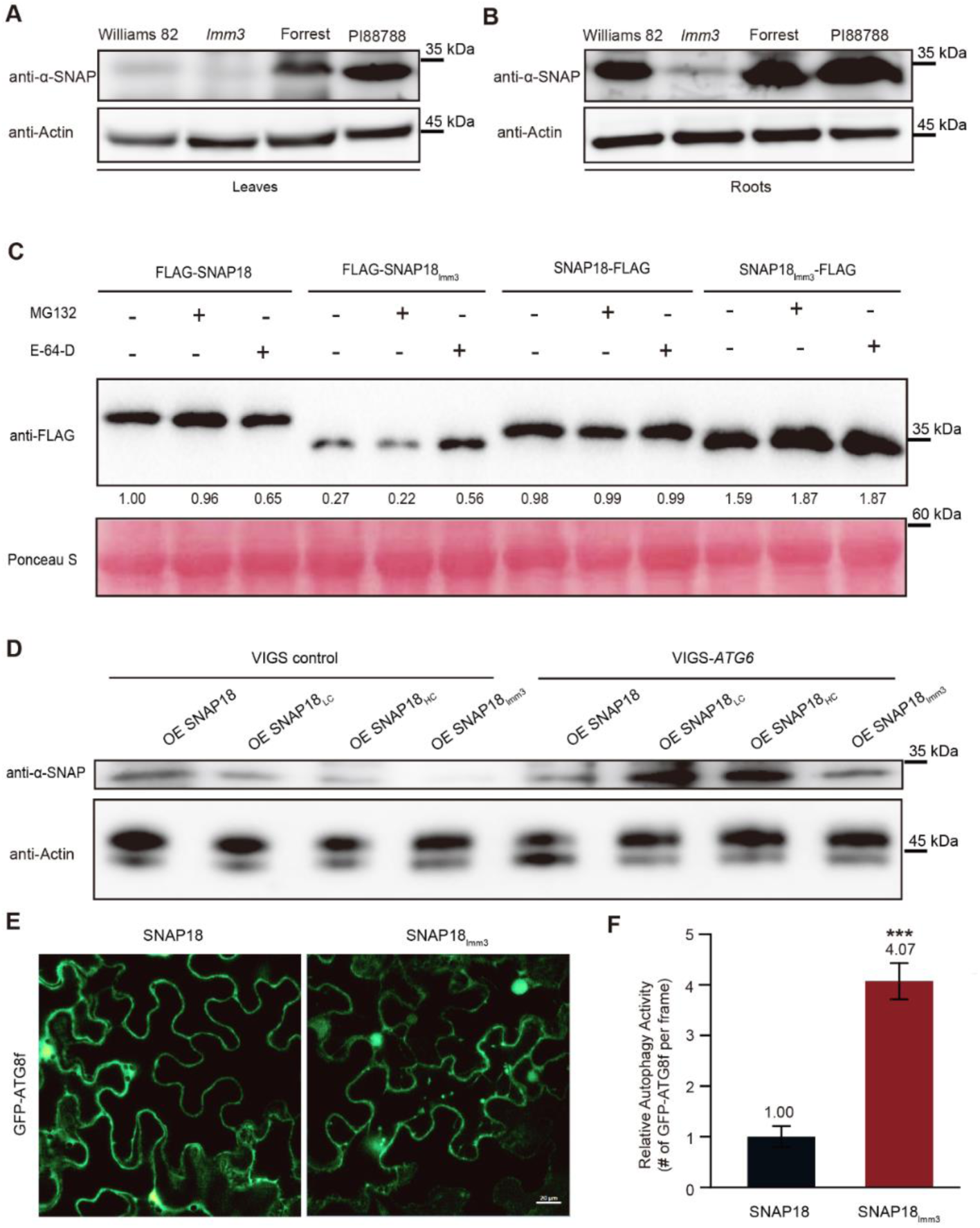
Soybean SNAP18lmm3 is degraded through autophagy. (A, B) Western blot with anti-α-SNAP antibody in leaves (A) and roots (B) from 7-d-old Williams 82, *lmm3*, Forrest and PI88788 plants. Actin was used as the loading control. (C) Western blot with anti-FLAG antibody of SNAP18 and SNAP18lmm3 abundance in transiently transgenic *N. benthamiana* leaves expressing FLAG*–*tagged SNAP18 and SNAP18lmm3 treated with E-64-D and MG132. Ponceau S staining was used as the loading control. The numbers between the anti-FLAG and Ponceau S panels represent the relative protein levels, normalized to lane 1, based on the grayscale ratio of anti-FLAG to Ponceau S. Lane 1 was set to ‘1’ for comparison purposes. (D) Western blot using anti-α-SNAP antibody in samples isolated from *N. benthamiana* leaves transiently expressing untagged SNAP18, SNAP18LC, SNAP18HC and SNAP18lmm3 with or without virus-induced *ATG6* gene silencing. Actin was used as the loading control. (E) Confocal microscopy of the autophagosome marker GFP–ATG8f transiently co-expressed with SNAP18 (left) and SNAP18lmm3 (right) in *N. benthamiana* leaves. Scale bar = 20 μm. (F) Quantification of GFP–ATG8f fluorescent aggregates representing autophagosomes per frame in (E). Values were normalized to GFP–ATG8f and are presented as means ±SEM (*** *P* < 0.001, Student’s *t*-test).

To validate this, we silenced the essential autophagy gene *ATG6* in *N. benthamiana* using virus-induced gene silencing (VIGS) vectored by tobacco rattle virus (TRV) (Liu et al. 2005). We also expressed *SNAP18*, *SNAP18_HC_*, *SNAP18_LC_*, and *SNAP18_lmm3_* from the *35S* promoter in separate spots within the same leaves, which were then treated with TRV alone or subjected to *ATG6* silencing. At 60 hours post-infection (hpi), SNAP18_lmm3_ abundance was restored upon *ATG6* silencing compared to the control treatment (Figure 4D). Furthermore, co-expression of GFP-ATG8f with native SNAP18 and SNAP18_lmm3_ resulted in more GFP-ATG8f-labeled autophagosomes in the presence of SNAP18_lmm3_ (Figure 4E,F). Collectively, these findings confirm the hypothesis that SNAP18_lmm3_ is degraded via autophagy.

### Constitutive Activation of Autophagic Flux Maintains Cellular Homeostasis in *lmm3*

We next aimed to clarify the extent of autophagy activity in the *lmm3* mutant using several experimental approaches. First, an ATG8/LC3 lipidation assay was employed (Luo and Zhuang 2018). ATG8 becomes lipidated with phosphatidylethanolamine (PE), resulting in the formation of an ATG8-PE complex, which heralds the initiation of phagophore formation, a precursor to autophagosome formation. We used uninoculated root segments from Williams 82, *lmm3*, Forrest, and PI88788. Following treatment with 100 μM E-64-D, western blotting was conducted using a custom anti-ATG8f antibody that detects both ATG8f and its lipidated form ATG8f-PE. Remarkably, we observed a substantial increase in ATG8f and ATG8f-PE in *lmm3* compared to wild-type Williams 82 and the other two *Rhg1*-resistant varieties (Figure 5A). Quantification of the ATG8f-PE:ATG8f ratio confirmed activation of autophagy in *lmm3* mutant at approximately 10.47 times higher levels than Williams 82 (Figure 5B). No statistically significant differences were found in the ATG8f-PE:ATG8f ratio between PI88788 and Williams 82 (Figure 5A,B). However, in the absence of E-64-D, the autophagic level in the *lmm3* mutant was comparable to that of Williams 82 (Figure S4). After 4 days of culture in sucrose-free medium under carbon starvation conditions, both Williams 82 and *lmm3* mutant showed a similar level of autophagic activation, indicating that the autophagy activation in *lmm3* is more likely associated with autophagic degradation rather than starvation-induced initiation (Figure S4). Second, root segments from Williams 82 and *lmm3* were subjected to transmission electron microscopy (TEM) analysis. This revealed conspicuous double-membraned autophagosomes in *lmm3* cells, whereas such structures were scarcely visible in the Williams 82 control (Figure 5C). Quantification of the number of autophagosomes per μm^2^ area indicated that *lmm3* had nearly nine times more autophagosomes than Williams 82, thereby confirming heightened autophagic activity in *lmm3* (Figure 5D). Third, we generated transgenic hairy roots in both Williams 82 and *lmm3* backgrounds expressing *35S_pro_:GFP-ATG8f* as a marker for autophagy and assessed localization by confocal microscopy. Speckled GFP-ATG8f fluorescence was prominent in the vacuoles of *lmm3* root cells, indicative of autophagosome formation. In contrast, GFP-ATG8f fluorescence in Williams 82 was uniformly distributed throughout the cell membrane and cytoplasm, without any noticeable aggregation (Figure 5E,F). Lastly, qRT-PCR analysis of several *ATG* marker genes at 8 DAG revealed that *lmm3* had up-regulated *ATG3*, *ATG6*, and *ATG8f* expression compared to the control (Figure S5). Collectively, these multi-dimensional lines of evidence confirm that root cells in the *lmm3* mutant sustain a significantly higher level of basal autophagy compared to wild-type and other resistant varieties.

**FIGURE 5.**
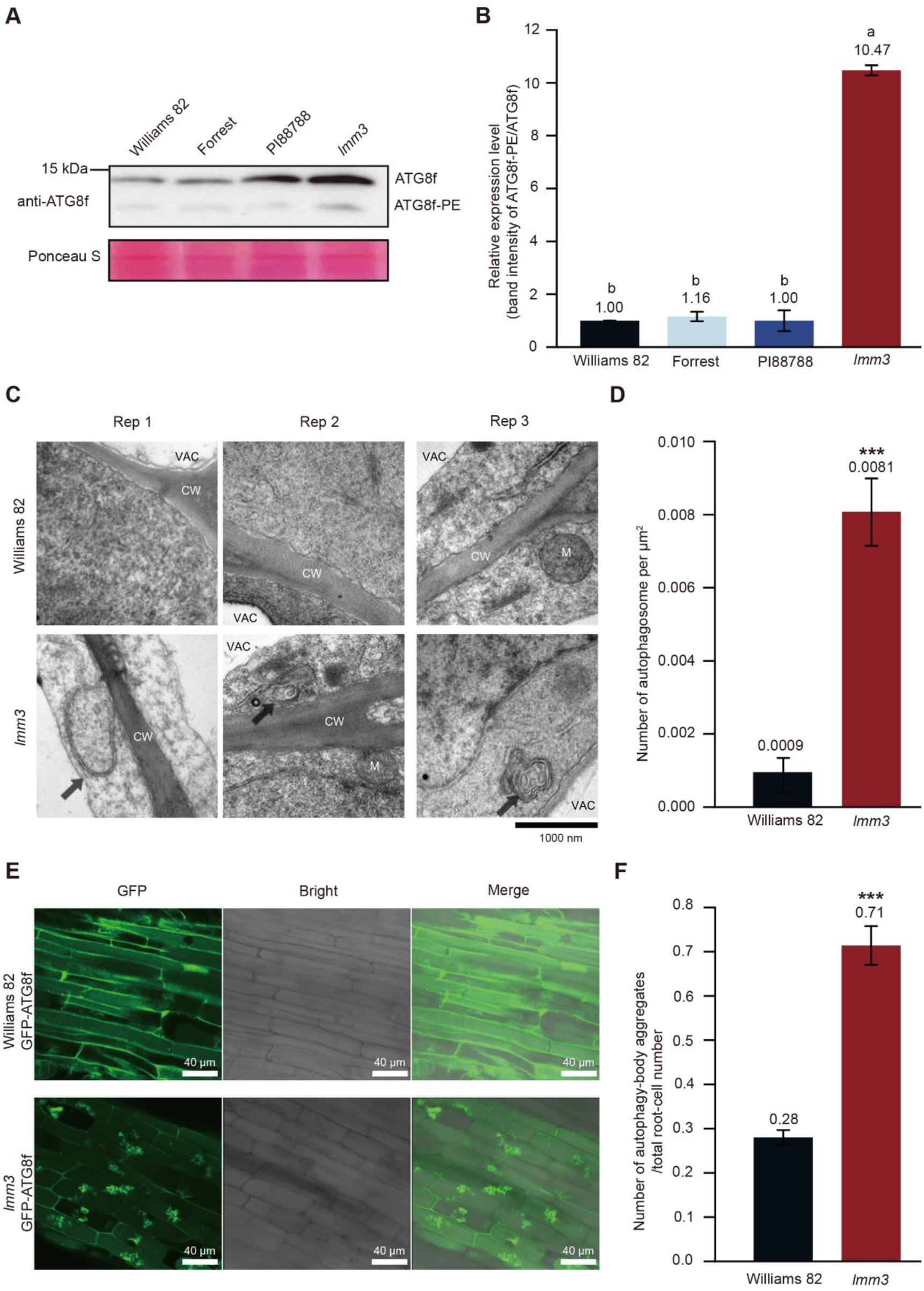
Autophagy is activated in the soybean *lmm3* mutant. (A) Western blot with anti-ATG8f antibody that detects PE modification in 7-d-old root extracts from Williams 82, Forrest, PI88788 and the *lmm3* mutant after treatment with 100 μM E-64-D. Ponceau S staining was used as the loading control. (B) Ratio of ATG8f*–*PE to ATG8f based on (A). Data are the mean ±SEM of n = 2. One-way ANOVA, different letters represent *P* < 0.05. (C) Transmission electron microscopy images to detect the presence of autophagosomal structures, highlighted by double-membrane autophagosomes (arrows) in root cells of wild-type Williams 82 and *lmm3* across three biological replicates. Cell wall (CW), mitochondrion (M), and vacuole (VAC) are labeled accordingly. Scale bar = 1 μm. (D) Quantification of autophagosomes per square micrometer of total cell image area as imaged in (C). Data are the mean ±SEM of n ≥ 40. \*\*\**P* < 0.001, Mann*–*Whitney test. (E) Confocal-microscopy analysis of transient GFP–ATG8f fluorescence as a marker of autophagy induction in roots of 7-d-old Williams 82 and *lmm3* seedlings. Scale bar = 40 μm. (F) Ratio of autophagy-body aggregates to the total root-cell number based on images picted in (E). Data are the mean ±SEM of n ≥ 23. \*\*\**P* < 0.001, Mann*–*Whitney test.

### Proteomic Analysis Reveals Basal Pre-Activation of the Autophagy Machinery in the *lmm3* Mutant

To further validate SNAP18_lmm3_’s role in autophagy regulation, we performed label-free quantitative proteomic profiling on Williams 82 and *lmm3* roots under both mock and SCN-infected (6 dpi) conditions. This confirmed up-regulation of several core autophagy (ATG) proteins, with ATG2 (Glyma.02G133400; 1.49-fold), ATG3 (Glyma.09G231000; 1.47-fold), and ATG12 (Glyma.07G038100; 1.45-fold) all up-regulated in *lmm3* roots following SCN infection (*lmm3*_SCN_vs_*lmm3*_Mock) (Table S1). Furthermore, another isoform of ATG3 (Glyma.12G005700; 1.47-fold) was higher in infected *lmm3* samples compared to infected wild-type (*lmm3*_SCN_vs_Wm82_SCN) (Table S1). Two ATG16 isoforms (Glyma.16G019300; 3.08-fold and Glyma.14G048000; 1.53-fold) and ATG27 (Glyma.11G233000; 1.45-fold) were all constitutively elevated in uninfected *lmm3* roots (*lmm3*_Mock_vs_Wm82_Mock) (Table S1). The constitutive elevation of these ATG proteins strongly suggests a basal pre-activation of the autophagy machinery in the *lmm3* mutant. This is consistent with the known resistance mechanism in plant-nematode interactions, where pre-activation of core defense components facilitates rapid responses to pathogen invasion (Yusuf and Bello 2025). These proteomic data also indicate that the *SNAP18_lmm3_* mutation confers resistance by reprogramming vesicular trafficking (notably the phagosome pathway) and pre-activating defense pathways. This pre-activation allows *lmm3* to mount an efficient and targeted response upon SCN infection, avoiding the broad, energy-intensive metabolic perturbations characteristic of the wild type (Figure S6). These experiments further support our hypothesis that autophagic activation in the *lmm3* mutant is associated with increased autophagic activity rather than starvation-induced initiation.

### NSF Acts as a Negative Regulator of SCN Resistance in Soybean

Based on our model that the disruption of the SNAP18-NSF interaction triggers resistance, we further elucidated a mechanism by which SNAP18_lmm3_ suppresses susceptibility to SCN, suggesting that NSF may act as a susceptibility factor in SCN challenge. To validate this, we performed an SCN demographic assay in transgenic hairy roots. The results revealed that nematode development was significantly restricted in NSF RNAi transgenic roots compared to control roots in the susceptible Williams 82 background. In contrast, no major differences in nematode development were observed between NSF RNAi and control roots in the resistant Forrest background (Figure S7). These findings underscore NSF’s role as a negative regulator of SCN resistance and suggest that the nematode relies on the host’s functional SNAP18/NSF machinery for successful parasitism.

### The *lmm3* Truncation Triggers a Competitive Binding Switch Between NSF and ATG8f

Using GFP-tagged SNAP18 as bait in an IP-MS assay in transformed soybean roots and leaves, we identified several ATG8 isoforms, including ATG8f (Glyma.17G140700) and other family members (Glyma.06G306300, Glyma.07G261000, Glyma.01G210200, and Glyma.02G008800), as potential binding partners (Table S2). BiFC and co-immunoprecipitation (co-IP) experiments confirmed interactions between SNAP18, as well as its mutant SNAP18_lmm3_, with ATG8f (Figure 6A,B). Notably, while ATG8f interacts with both isoforms, only SNAP18_lmm3_ undergoes degradation. We hypothesized that NSF, as a ligand for SNAP18, may suppress the SNAP18-ATG8 interaction. To verify this, we co-expressed SNAP18, the N-terminal portion of NSF, and ATG8f in *N. benthamiana* for co-IP (Figure 6C). *In vitro* pull-down experiments demonstrated that GST-SNAP18_lmm3_ effectively pulled down 6xHis-ATG8f in the presence of 6xHis-NSF_1-205_ (Figure 6D). However, the presence of NSF_1-205_ strongly suppressed the interaction between SNAP18 and ATG8f (Figure 6D). Additionally, yeast two-hybrid experiments showed that the addition of NSF_1-205_ significantly reduced the interaction between BD-SNAP18 and AD-ATG8f, whereas it had no effect on the interaction between BD-SNAP18_lmm3_ and AD-ATG8f (Figure 6E). Together, these results define a competitive binding switch between NSF and ATG8f, where the *lmm3* truncation shifts the fate of SNAP18 from NSF-mediated stabilization to autophagic degradation.

**FIGURE 6.**
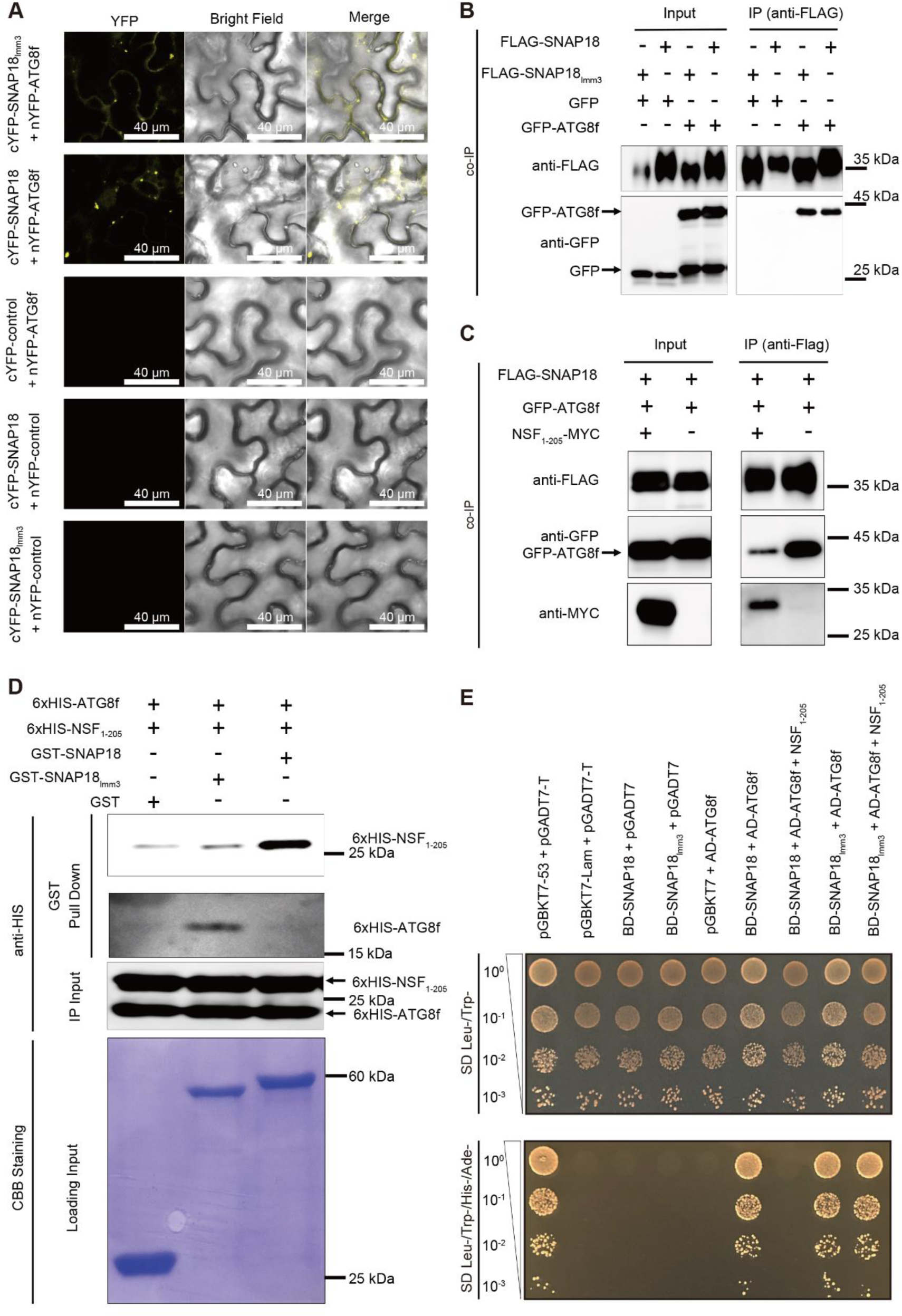
SNAP18lmm3 preferentially interacts with ATG8 over NSF’s N-terminal domain. (A) Representative images from BiFC assays of interactions between SNAP18lmm3 and ATG8f. nYFP*–*ATG8f was transiently co-expressed with N-terminal cYFP-tagged SNAP18 and SNAP18lmm3 in *N. benthamiana* leaves. Negative controls with different pairings are as indicated. Scale bars = 40 μm. (B) Co-immunoprecipitation analysis of FLAG*–*SNAP18 and FLAG*–*SNAP18lmm3 interactions with GFP*–*ATG8f. GFP*–*ATG8f was detected in both the immunoprecipitants of FLAG*–*SNAP18lmm3 and FLAG*–*SNAP18. (C) Co-immunoprecipitation of FLAG*–*SNAP18, GFP*–*ATG8f and NSF1-205*–*MYC. The interaction between FLAG*–*SNAP18 and GFP*–*ATG8f is suppressed by NSF1–205*–*MYC, where NSF1–205 refers to the N-terminal 205 amino acids of NSF. (D) Pull-down analysis of 6 ×His*–*ATG8f and 6 ×His*–*NSF1-205 with GST-tagged SNAP18 and SNAP18lmm3. Only GST*–*SNAP18lmm3 successfully pulled down 6 ×His*–*ATG8f. (E) Yeast two-hybrid analysis of the interaction between BD*–*SNAP18 or BD*–*SNAP18lmm3 and AD*–*ATG8f, with or without NSF1-205. Yeast transformants were cultured on selective medium (SD Leu-/Trp-/His-/Ade-). The addition of NSF1-205 affected the interaction between BD*–*SNAP18 and AD*–*ATG8f, but did not affect the interaction between BD*–*SNAP18lmm3 and AD*–*ATG8f. pGBKT7-53 + pGADT7-T was used as a positive control, and pGBKT7-Lam + pGADT7-T was used as a negative control.

### Structural Basis and Spatial Regulation of the “Self-Degrading Toxin” Model

To further investigate these differential interactions, we used AlphaFold3 to predict the interaction structures of SNAP18 with NSF and ATG8f, respectively, and aligned the interaction residues of SNAP(3J97)-NSF(3J97) [PDB ID code 3J97] with those from the predicted SNAP18-NSF structure (Figure S8 and Figure S9A,B). Key residues in the AlphaFold3-predicted SNAP18-NSF structure align closely with those in the binding interface of SNAP(3J97)-NSF(3J97). This alignment confirms the credibility of the AlphaFold3-predicted structure (Figure S8; Figure S9A and Table S3).

Both NSF and ATG8f interact with the C-terminus of SNAP18 as suggested by AlphaFold3 structure-interaction prediction. The predicted SNAP18-NSF structure has lower binding energy (−1.2 kcal/mol) compared to the predicted SNAP18-ATG8f structure (0.8 kcal/mol). Additionally, the interaction surface of SNAP18-NSF (608.2 Å²) is larger than that of SNAP18-ATG8f (413.8 Å²) and includes more direct interaction residues with shorter interaction distances (Figure S9A,B; Table S3). These findings suggest that SNAP18 preferentially binds to NSF, which competitively occupies its C-terminus, thereby preventing interaction with ATG8f and highlighting the potential importance of this site in autophagy regulation (Figure S9C). These results support a model in which ATG8f binds both isoforms, but only SNAP18_lmm3_ is routed to autophagy for degradation, consistent with NSF competitively limiting ATG8 access to the SNAP18 C terminus.

## Discussion

### A Competitive Molecular Switch Determines the Biochemical Fate of SNAP18_lmm3_

In this study, we characterized a novel autophagic degradation mechanism that orchestrates the homeostasis of the truncated SNAP18 isoform (SNAP18_lmm3_), thereby reconciling robust soybean resistance to *Heterodera glycines* with sustained plant growth. Our findings reveal that SNAP18_lmm3_ acts as a “self-degrading toxin,” whose biological fate is determined by a competitive molecular switch between vesicular trafficking and autophagic clearance. This mechanism establishes a new cellular paradigm for understanding how plants manage potentially cytotoxic immune components to ensure systemic survival while mounting effective pathogen defense. The C-terminal helical bundle of α-SNAP is evolutionarily conserved across eukaryotes, serving as the primary interface for binding the NSF hexamer—an interaction essential for the disassembly and recycling of SNARE complexes during vesicle docking and fusion (Bombardier and Munson 2015). In wild-type soybean, the high-affinity association between SNAP18 and NSF effectively masks the ATG8-interacting motifs (AIMs) or structural surfaces required for autophagic recognition. However, the 24-amino-acid truncation in *lmm3* disrupts this canonical interaction, generating a population of “orphaned” SNAP18 proteins that are excluded from vesicular trafficking (Bayless et al. 2016; Bayless et al. 2018). Our results demonstrate that this interaction deficit triggers a functional shift: reorientation, in the absence of the “protective shield” provided by NSF, SNAP18_lmm3_ becomes an unimpeded target for ATG8f-mediated autophagy. This ligand-dependent fate determination represents a sophisticated protein quality control mechanism, wherein the disruption of a critical trafficking complex is immediately sensed and mitigated by the autophagic machinery.

Notably, the *α-SNAP* gene family in soybean comprises five members, with several variants (e.g., *SNAP11*, *SNAP02*) in resistant accessions also harboring C-terminal truncations linked to SCN resistance (Basnet et al. 2022; Lakhssassi et al. 2017; Matsye et al. 2012; Shaibu et al. 2022; Usovsky et al. 2023) (Figure S10). These mutations, alongside previously reported mis-spliced mRNAs that impair NSF binding, may similarly disrupt syncytial function and engage autophagy as a conserved defense mechanism across diverse α-SNAP loci. This suggests that the competitive switch between NSF and ATG8f binding could represent a generalizable model for α-SNAP-mediated resistance, expanding beyond the specific SNAP18_lmm3_ variant characterized here.

### The “Self-Degrading Toxin” Model Balances Cellular Homeostasis and Immunity

A defining feature of SNAP18_lmm3_ is its ability to confer dominant SCN resistance without the systemic yield penalties typically associated with autoimmune or lesion-mimic mutants. We propose a dual-mode “self-degrading toxin” model to explain this balance (Figure 7), wherein autophagy acts as a systemic safety valve to preserve plant vigor while enabling targeted pathogen arrest:

1. **Systemic Detoxification in Non-Infected Tissues:** In non-infected tissues, the inherent cytotoxicity of SNAP18_lmm3_, stemming from its interference with normal membrane fusion is kept in check by a constitutively activated autophagic flux (Tang and Bassham 2018). Our proteomic data, showing the basal pre-activation of core ATG proteins (e.g., ATG2, ATG16), suggest that the *lmm3* mutant maintains a “primed” state of cellular homeostasis to clear aberrant protein aggregates.
2. **Localized Pathogen Arrest Upon SCN Infection:** Upon SCN infection, SNAP18_lmm3_ hyper-accumulates specifically within the multinucleated syncytia, reaching concentrations nearly four times higher than in adjacent cells (Bayless et al. 2016; Bayless et al. 2018). This rapid, localized accumulation likely overwhelms the local autophagic capacity, leading to the collapse of vesicular trafficking, targeted cell death, and ultimately the arrest of nematode development. Critically, this response is spatially restricted to the syncytium, avoiding the broad metabolic perturbations that often accompany systemic immune activation.

**FIGURE 7.**
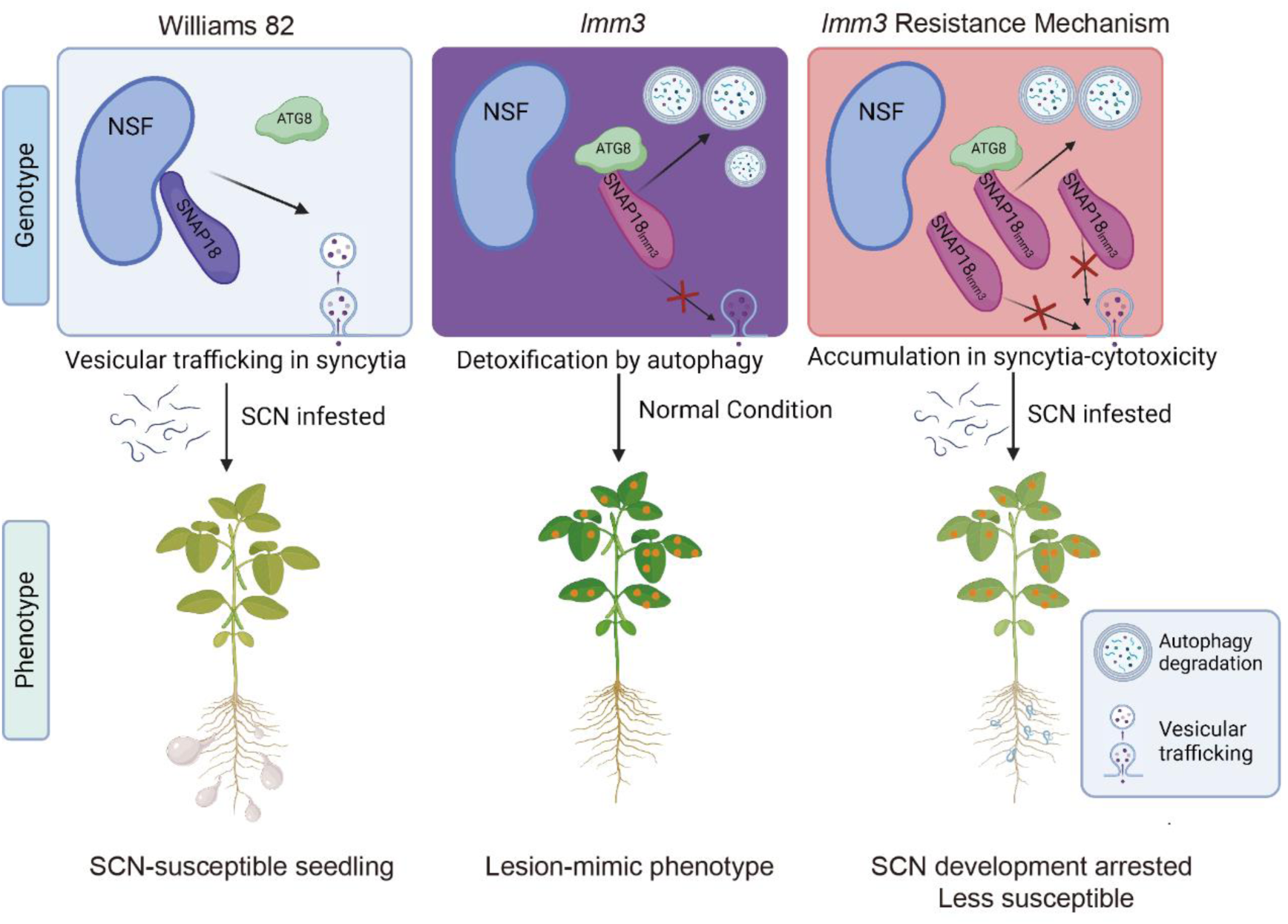
Working model of the “self-degrading toxin” mechanism balancing vesicular trafficking and autophagy in soybean SCN resistance. Proposed model of molecular and phenotypic mechanisms of α-SNAP function in soybean resistance to SCN. Left: In the wild type, SNAP18 binds NSF, supporting vesicle trafficking and allowing SCN cyst formation. Middle: In the *lmm3* mutant, truncated SNAP18lmm3 fails to bind NSF and is degraded by autophagy, causing lesion-mimic symptoms. Right: Under SCN infection, toxic SNAP18lmm3 accumulates in syncytia, causing cell death and SCN resistance. Symbols indicate autophagy degradation and vesicle trafficking. Created with BioRender. Han, S. (2025). Retrieved from https://biorender.com/y40y278.

This model addresses a longstanding challenge in plant pathology: how to achieve robust resistance without compromising growth. By coupling the cytotoxic potential of a trafficking-disrupting protein with autophagic regulation, soybean has evolved a strategy to deploy defense at the site of infection while minimizing fitness costs elsewhere. This stands in contrast to traditional *Rhg1*-mediated resistance, which relies on gene copy-number expansion and often requires compensatory NSF variants (e.g., NSF_RAN07_) to mitigate α-SNAP-induced toxicity (Bayless et al. 2018).

### NSF Functions as a Host Susceptibility Factor for SCN Parasitism

Our discovery that NSF RNAi restricts nematode development in susceptible cultivar (Williams 82) confirms NSF as a negative regulator of SCN resistance (Figure S7). This finding aligns with the emerging paradigm that pathogens often hijack essential host cellular processes to establish infection, and disrupting these “susceptibility (S) factors” can confer broad-spectrum resistance (Bent 2022). Specifically, the successful formation of syncytia, a high-metabolism nutrient sink, requires an exceptionally efficient SNARE-recycling machinery driven by the SNAP18-NSF complex. By interfering with this cycle, SNAP18_lmm3_ directly targets a metabolic vulnerability of the nematode, highlighting the potential of targeting host S factors as a complementary strategy to conventional R-gene-based resistance.

Notably, NSF RNAi had no significant effect on nematode development in the resistant Forrest background, suggesting that *rhg1-a*-mediated resistance already impairs the SNAP18-NSF pathway. This further supports our model that disruption of the SNAP18-NSF interaction is a central driver of SCN resistance, with SNAP18_lmm3_ representing a naturally occurring variant that achieves this disruption in a single-copy *rhg1-c* background.

### Biotechnological Implications and Future Directions

From a biotechnological perspective, the single-copy nature of SNAP18_lmm3_ and its “self-regulating” behavior make it an ideal candidate for precision genome editing. Recreating this specific 24-amino-acid truncation in high-yielding elite cultivars could bypass the need for traditional marker-assisted backcrossing and avoid the linkage drag associated with the *Rhg1* genomic block. Additionally, stacking *SNAP18_lmm3_* with other resistance loci (e.g., *Rhg4*) may enhance resistance durability by targeting multiple stages of SCN infection.

Several critical questions remain to be addressed. First, the specific signals that trigger the hyper-accumulation of SNAP18_lmm3_ within syncytia, whether through enhanced translation, localized protein stabilization, or reduced autophagic activity at feeding sites, require further investigation. Second, the role of autophagy in degrading nematode effectors or other host defense suppressors warrants exploration, as autophagy may contribute to resistance through both toxin clearance and direct pathogen targeting. Third, evaluating the stability of this autophagy-mediated balance across diverse environmental conditions (e.g., nutrient limitation, drought) will be critical for translating this finding into field-applicable resistance.

Finally, our work highlights the broader importance of autophagy in balancing crop growth and immunity. While autophagy has been implicated in plant responses to various stresses, its role as a regulatory hub for trafficking-related cytotoxicity and resistance to parasitic nematodes is poorly understood. By elucidating this mechanism, we provide a novel framework for engineering durable crop resistance through the modulation of cellular homeostasis, an approach that could be extended to other plant-pathogen interactions where vesicular trafficking or protein toxicity plays a central role.

In summary, our study uncovers a competitive molecular switch between NSF and ATG8f binding that governs SNAP18 function, establishes the “self-degrading toxin” model for balancing growth and immunity, and identifies NSF as a key susceptibility factor for SCN. These findings advance our understanding of plant-nematode interactions and offer new strategies for the genetic improvement of disease-resistant crops.

## Materials and Methods

### Plant Materials and Growth Conditions

The soybean cultivars Williams 82, Forrest (*rhg1-a* genotype) and PI88788 (*rhg1-b* genotype) were utilized in this study. For controlled-environment experiments, soybean plants were cultivated in a climate chamber maintained at 26 ℃, 50% relative humidity under a 14-hour photoperiod (560 μmol photons m^-2^ s^-1^). These chamber-based assays, including seedling-stage SCN inoculation and growth phenotypic characterization, were conducted to complement field results by minimizing environmental variability and ensuring high reproducibility.

For transient expression assays, *Nicotiana benthamiana* seeds were initially germinated on moist filter paper at 25 ℃ in the dark for 3 d. Following germination, seedlings were transplanted into pots containing a 2:1 mixture of soil and vermiculite and cultivated in a growth chamber set to 24 ℃, 60% humidity with a 12-hour light/dark cycle with a light intensity of 270 μmol photons m^-2^ s^-1^. All transient expression experiments were performed using four-week-old plants at the appropriate physiological stage.

### Nematode-Infection Assays

Nematode cultures HG Type 0 (Race 3) were maintained and used according to established protocols with slight modifications (He et al. 2025). Hatched second-stage juveniles (J2s) obtained from crushed cysts were collected for inoculation. A suspension containing 1,000 J2s in 1 mL was inoculated onto roots of 10-day-old plants at 26 °C. The use of J2s at a standardized density allows precise control of infection under uniform conditions, which is suitable for quantifying early infection events. In contrast, chamber screenings were conducted with cysts/eggs at higher inoculum densities to ensure robust infection establishment under variable soil and environmental conditions. The number of nematodes inoculated at different stages was counted, and fuchsin-stained roots were observed using an Olympus BX53 microscope (Olympus, Japan). Each experiment included more than 10 plants per treatment and was repeated at least three times.

### Plasmid Construction

For transient-expression vectors, open reading frames (ORFs) of soybean *ATG8f* (*Glyma.17G140700*), *SNAP18*, *SNAP18_LC_*, *SNAP18_HC_* and *SNAP18_lmm3_* were amplified from cDNA synthesized from Williams 82, Forrest, PI88788 and *lmm3* using the HiScript III 1st Strand cDNA Synthesis Kit (Vazyme) and KOD-Plus-High fidelity DNA polymerase (Toyobo). Each ORF was assembled with the double CaMV *35S* promoter with the TMV omega enhancer (pICH51288) and the nopaline synthase (*NOS*) terminator in the binary vector pAGM4673 (MoClo Tool Kit). The assembly process was carried out using Golden Gate cloning (Weber et al. 2011).

For the preparation of BiFC vectors, expression constructs for nYFP or cYFP fusions were generated using the CaMV *35S* promoter, following a similar protocol as described previously (Zhao et al. 2013). The ORFs encoding *SNAP18s* with stop codon, *ATG8f* with stop codon, or *NSF* without stop codon were amplified by overhang PCR. ORFs were flanked by a specific LIC1 adaptor (5’-CGACGACAAGACCGTGACC-3’) at the 5’ end and LIC2 adaptor (5’-GAGGAGAAGAGCCGT-3’) at the 3’ end. PCR products were then purified using the QIAquick Gel Extraction Kit (Qiagen). Ligase Independent Cloning (LIC), as described previously (Weber et al. 2011), was employed to fuse the N-terminal cYFP with SNAP18s, N-terminal nYFP with ATG8f and to fuse the C-terminal nYFP with NSF.

To generate co-IP vectors, full-length *ATG8f* fused with N-terminal *GFP* (using a *ATG8f* cDNA with a stop codon) was cloned using LIC (Du et al. 2013) into the binary expression vector pJG045, driven by the CaMV *35S* promoter. Similarly, a construct containing C-terminal *SNAP18s* and N-terminal 3xFLAG tag was also cloned into pJG045 using LIC. The primers used in this study are listed in Table S4.

### Transient Expression in *Nicotiana benthamiana*

*N. benthamiana* plants with 2-3 fully expanded leaves were used for infiltration using *Agrobacterium tumefaciens* GV3101 (pMP90) following a previously described protocol (Han et al. 2023). For secGFP co-expression experiments, secGFP was co-infiltrated at an OD_600 nm_ of 0.1 along with either a specified SNAP18 construct or an empty-vector control, all at an OD_600 nm_ = 0.6.

### Co-Immunoprecipitation (co-IP)

Four-week-old fully expanded *N. benthamiana* leaves were selected and infiltration was performed by applying a bacterial solution at OD_600 nm_ = 0.6. Expression was allowed to proceed for 60 h. Four different plants were used, and approximately 2 g of leaf tissue per treatment was collected to constitute one biological replicate. Tissue was immediately frozen using liquid nitrogen and manually ground in a pre-chilled mortar and pestle. Subsequently, 4 mL of protein extraction buffer (50 mM Tris·HCl pH 7.5, 150 mM NaCl, 5 mM EDTA, 0.2% [v/v] Triton X-100, 10% [v/v] glycerol, and 1/100 Sigma protease inhibitor cocktail) was added and the sample further homogenized. Lysates were transferred to tubes and centrifuged at 6,000 *g* for 10 min at 4°C, repeated three times, to remove insoluble debris. The supernatant was incubated with prewashed anti-FLAG Nanobody Magarose beads (AlpaLifeBio) for 3 h at 4°C. The bead precipitates were then washed four times with ice-cold immunoprecipitation buffer at 4°C.

Protein concentrations were determined using Bradford assay. Samples were subsequently subjected to analysis by western blot using anti-FLAG (Sigma, 1:10,000), anti-GFP (Sigma, 1:10,000) or anti-MYC antibodies (Cell Signaling Technology, 1:10,000). Chemiluminescence detection was carried out with SuperSignal Dura chemiluminescent substrate (Thermo Scientific) and visualized using a ChemiDoc MP chemiluminescent imager (Bio-Rad).

### Immunoprecipitation-Mass Spectrometry (IP-MS)

To identify SNAP18-interacting proteins, GFP-tagged SNAP18, derived from the roots and leaves of transgenic *lmm3 LMM3_pro_:LMM3-GFP* complementation lines, was used as bait for immunoprecipitation. Protein complexes were immunoprecipitated using GFP-Trap (ChromoTek gtma) and subjected to SDS-PAGE, followed by in-gel digestion.

For protein digestion, samples were reduced with 5 mM DTT at 37°C for 60 min, alkylated with 10 mM IAA in the dark for 45 min, and digested overnight at 37°C with trypsin (1:50 enzyme-to-protein ratio) in 50 mM ammonium bicarbonate. Peptides were acidified with 1% formic acid (FA), desalted using C18 columns, and eluted with 100% acetonitrile (ACN) before being vacuum-dried.

Peptides were analyzed by nanoflow LC-MS/MS using a Q Exactive HF-X Orbitrap mass spectrometer (Thermo Fisher Scientific) coupled to an EASY-nLC 1200 system. Peptides were loaded onto a 25 cm C18 column and separated using a 60-minute gradient from 8% to 95% solvent B (80% ACN, 0.1% FA) at a flow rate of 600 nL/min. The mass spectrometer operated in data-dependent mode, acquiring MS spectra at 120,000 resolution, followed by MS/MS of the top 40 precursors with higher-energy collisional dissociation (HCD) at 27% normalized collision energy.

Raw data were analyzed using Proteome Discoverer with the Sequest HT search engine against the UniProt soybean proteome database. Search parameters included trypsin specificity (up to two missed cleavages), carbamidomethylation of cysteines (fixed modification), and oxidation of methionine (variable modification). A 15 ppm precursor mass tolerance and a 0.02 Da fragment mass tolerance were applied. Peptide spectral matches were filtered to a false discovery rate (FDR) <1% using Percolator, and protein identification was based on at least one unique peptide.

### BiFC Assay

To investigate protein-protein interactions, BiFC assays were performed as previously described (Zhao et al. 2013). *A. tumefaciens* GV3101 carrying the indicated constructs fused to split YFP fragments (nYFP and cYFP) was co-infiltrated into *N. benthamiana* leaves. Two days post-infiltration, the infiltrated leaves were observed under a confocal microscope (Zeiss 880) to detect YFP fluorescence. Imaging settings were kept consistent to ensure reproducibility, and excitation/emission wavelengths of 514/519-587 nm were used to visualize YFP signals.

### Pull-Down Assay

To assess protein interactions, pull-down assays were conducted as previously described (Han et al. 2023). GST-tagged bait proteins were expressed and purified from *E. coli* BL21 (DE3) cells using glutathione-Sepharose 4B beads (GE Healthcare). About 1 mg bait and prey proteins were incubated in pull-down buffer A (50 mM Tris-Cl, pH 7.5, 100 mM NaCl, 0.1 mM EDTA, 0.1 mM EGTA, 0.2% Triton X-100, 0.1% β-mercaptoethanol, 1 mM PMSF, and complete protease inhibitor) for 3 h at 4 ℃ with gentle agitation. The beads were washed five times with buffer B (50 mM HEPES, pH 7.5, 100 mM NaCl, 0.1 mM EDTA, 1 mM PMSF, and complete protease inhibitor), and bound proteins were eluted by boiling with 2×SDS loading buffer for 5 min. Proteins were analyzed by SDS-PAGE, followed by immunoblotting with anti-His antibodies (ABclonal).

### Confocal Microscopy

Confocal microscopy was conducted using an inverted Carl Zeiss laser-scanning confocal microscope (LSM 880) equipped with a 20× or 40× water-immersion objectives. Excised *N. benthamiana* leaves, infiltrated with *A. tumefaciens*, were observed approximately 72 h after infiltration. GFP was excited at 488 nm and the emitted fluorescence was detected using a 493-594 nm emission filter. For mCherry, excitation was at 561 nm, with fluorescence detected using a 580-630 nm emission filter. YFP was excited at 514 nm, and the emitted fluorescence was detected using a 520-560 nm emission filter.

For BiFC assays, YFP signal was acquired at 514 nm and a 519-620 nm range emission filter. Image acquisition was performed using a standardized scan area of 442.2 ×442.2 μm (for the 20× objective) or 212.55 × 212.55 μm (for the 40× water immersion objective) with a frame size of 1024 × 1024 pixels. The detector master gain setting ranged from 700 to 800 depending on signal intensity. A pinhole size of 1.01 AU was used for all *N. benthamiana* samples and 2.36 AU for soybean root samples. A minimum of 36 images were analyzed for each expression treatment across three independent experiments.

### Conventional Transmission Electron Microscopy (TEM)

Soybean root samples were harvested 14 d after transformation. Root segments in the elongation zone, approximately 2 mm long, were subjected to vacuum infiltration with 2.5% v/v glutaraldehyde. Following this, the samples were stained with 1% w/v osmium tetroxide for 4 h. Five rinses with 100 mM phosphate buffer (pH 7.0) were performed, with each rinse having a 30-min interval. The samples were then dehydrated using a graded series of ethanol (50%, 70%, 80%, 90%, 95%, and 100%) and pure acetone, each for 40 min. Subsequently, samples were embedded in Epon 812 resin (SPI Supplies Inc., USA). Sections with a thickness of 100 nm were prepared using a Leica UC 6 microtome and placed onto 100 mesh carbon-covered copper grids. Prior to examination, grids were stained with 2% v/v uranyl acetate and 2% v/v lead citrate for 15 min each. Examination and photography were performed using an H-7650 transmission electron microscope (Hitachi, Japan) operating at 80 kV, equipped with a Gatan 830 CCD camera (Gatan, USA). Representative images were captured from four independent root segments.

### Electron Microscopy and Immunolabeling

Electron microscopy and immunolabeling techniques were employed following a similar protocol described previously (Han et al. 2023). Root segments from SCN-inoculated plants were manually sectioned into approximately 2-mm long pieces. These sections were then subjected to vacuum infiltration in fixation buffer consisting of 0.1% v/v glutaraldehyde and 4% v/v paraformaldehyde in 0.1M sodium phosphate buffer at pH 7.4, followed by overnight incubation. The samples were subsequently dehydrated using a series of ethanol concentrations (50%, 70%, 90%, 95%, and 100%) and embedded in LR White resin. Longitudinal sections of about 90-nm thickness were obtained using an ultramicrotome (UC-6; Leica). For immunogold labeling, the sections were mounted onto nickel slot grids.

To prepare grids for labeling, they were first activated by incubating in drops of 50 mM glycine/PBS for 15 min, followed by blocking in drops of blocking solution for goat gold conjugates (Aurion) for 30 min. Grids were then equilibrated in incubation buffer (0.1% w/v BSA-C/PBS). Next, grids were incubated with anti-SNAP18 antibodies diluted 1:1,000 in incubation buffer overnight at 4°C. After washing five times in incubation buffer, grids were incubated with goat anti-rabbit secondary antibody conjugated to 15-nm gold (Aurion) diluted 1:50 in incubation buffer for 2 h. Following six washes in incubation buffer and two washes in PBS, grids were fixed using 2% v/v glutaraldehyde in 0.1 M phosphate buffer for 5 min. Finally, grids were further washed in phosphate buffer and water. Images were captured using a MegaView III digital camera mounted on a Philips CM120 transmission electron microscope.

### Antibody Production

Custom affinity-purified polyclonal antibodies against soybean -SNAP18s and ATG8f were generated by ABclonal. Antibodies against SNAP18s were raised in rabbits using a synthetic peptide ‘KAEKKLSGWGLFGSK’ corresponding to residues 14-28 near the N-terminus of SNAP18s. Antibodies against ATG8f were raised in rabbits using a synthetic peptide ‘YSGENTFGDLTSH’ corresponding to residues 115-128 near the C-terminus.

### Western Blotting

Soybean root samples were rapidly frozen in liquid nitrogen and protein was extracted in buffer composed of 50 mM Tris·HCl pH 7.5, 150 mM NaCl, 5 mM EDTA, 0.2% v/v Triton X-100, 10% (v/v) glycerol, and a protease inhibitor mixture (Sigma, P9599). Extraction was performed by homogenizing samples in a PowerLyzer 24 (MO BIO) at 2,000 rpm for 3 cycles with a 15-sec interval. Protein was quantified by Bradford assay. Proteins (20 µg per lane) were separated on 12% SDS-PAGE gels and transferred onto PVDF membranes (Millipore) using a wet transfer system at 400 mA for 1 h in transfer buffer (25 mM Tris, 192 mM glycine, 20% (v/v) methanol). Membranes were blocked in 5% (w/v) non-fat milk in TBS-T buffer (50 mM Tris, 150 mM NaCl, 0.05% v/v Tween 20) for 1 hour before incubation with primary antibodies. Membranes were incubated overnight at 4°C with anti-SNAP18 antibody (1:1,000) in 5% (w/v) non-fat dry milk TBS-T. After four washes with TBS-T, horseradish peroxidase-conjugated goat anti-rabbit secondary antibody was added at 1:10,000 and incubated for 1 h at room temperature with gentle agitation. Membranes were then washed four times with TBS-T and subjected to chemiluminescence detection using SuperSignal West Pico or Dura chemiluminescent substrate (Thermo Scientific). Finally, membranes were imaged using a ChemiDoc MP chemiluminescent imager (Bio-Rad). Band intensities of target proteins from Western blots were quantified using ImageJ software. High-resolution images of the blots were imported into ImageJ, and the rectangular selection tool was used to outline and measure the pixel intensity of each protein band. Background correction was performed by subtracting the mean intensity of an adjacent, unstained region from each band measurement to ensure accurate quantification. Normalization was achieved by calculating the ratio of the target protein intensity to either an internal reference protein (e.g., β-actin) or the total protein staining intensity (Ponceau S). The normalized intensity was expressed as: Normalized Intensity = Target Protein Intensity / Reference Intensity. Quantifications were carried out across at least two independent biological replicates. Error bars represent the standard deviation (SD) or the standard error of the mean (SEM) calculated from these replicates.

### Yeast Two-Hybrid Assay

To construct the fusion protein expression vectors, the full-length coding sequences of SNAP18 and SNAP18_lmm3_ were subcloned into the pGBKT7 vector (Clontech), fused with the GAL4 DNA binding domain (BD) to generate the pGBKT7-Bait vectors. The full-length coding sequence of ATG8f was subcloned into the pGADT7 vector (Clontech), fused with the GAL4 transcriptional activation domain (AD) to generate the pGADT7-Prey vector. Additionally, the N-terminal sequence of NSF (amino acids 1-205) was subcloned into the pGADT7 vector without the activation domain (AD) to produce NSF_1-205_, which was used to assess its competitive effect on ATG8f binding. Yeast transformation was performed using PEG and lithium acetate buffer. BD-SNAP18 or BD-SNAP18_lmm3_ were co-transformed with AD-ATG8f, with or without the NSF_1-205_ plasmid, into the AH109 yeast strain and plated onto selective medium lacking Trp and Leu. Protein-protein interactions were assessed by the growth of yeast colonies on medium lacking Leu, Trp, His, and Ade. pGBKT7-53 and pGADT7-T were used as positive controls, while pGBKT7-Lam and pGADT7-T were used as negative controls.

### Generation of Transgenic Hairy Roots

Transgenic hairy roots were generated using *A. rhizogenes* ArQua1 as described previously (Melito et al. 2010). The cassette to over-express GFP-ATG8f in *N. benthamiana*, as described above, was cloned together with the GFP cassette into pAGM4673 using Golden Gate assembly. To generate hairy roots, cotyledons were transformed with either the GFP empty vector or the GFP-ATG8f construct following established protocols (Chen et al. 2024). Transgenic lines were screened for GFP expression under a fluorescence microscope to confirm successful transformation.

### Protein-Protein Interaction Structure Prediction

Protein sequences were submitted to the AlphaFold3 online platform (https://alphafoldserver.com/) for structural prediction. The resulting structure were then converted into **.pdb** files and uploaded to the online tool at https://www.ebi.ac.uk/msd-srv/prot_int/pistart.html to predict and evaluate the interaction interfaces. Finally, PyMOL (version 2.5.5) was used to visualize the predicted protein interaction structures.

### Amino-Acid Alignments

The amino-acid sequences of SNAP18, SNAP02, and NSF were obtained from Phytozome (https://phytozome-next.jgi.doe.gov/). SNAP(3J97) and NSF(3J97) correspond to GenBank accession numbers NP_542152 and XP_007606206, respectively. The amino-acid sequences of SNAP11_A240*_ and SNAP11_E244*_, found in SCN-resistant soybean varieties Forrest and Peking, were obtained from Lakhssassi et al. (Lakhssassi et al. 2017). Additionally, the sequence of SNAP02del, originating from PI 437654, was obtained from Usovsky et al. (Usovsky et al. 2023). Alignments were performed using the Clustal W algorithm in MEGA software (version 5.05) and visualized with GeneDoc. Alternatively, alignments were conducted using the L-INS-i (accurate) option in MAFFT software and visualized with CLC Sequence Viewer 8.0.

### Statistical analysis

All experiments were conducted with at least two independent biological replicates. Error bars in the figures represent either the standard deviation (SD) or the standard error of the mean (SEM), as indicated in each figure legend. Statistical analyses were performed using GraphPad Prism software (version 10.2.0).

To assess statistical significance, appropriate tests were selected based on data distribution and experimental design. These included two-tailed Student’s *t*-tests for normally distributed data, Mann-Whitney tests for non-parametric comparisons, and one-way or two-way ANOVA for multi-group analyses. A *P*-value < 0.05 was considered statistically significant. Statistical significance is denoted as follows: Asterisks indicate differences from two-tailed Student’s *t*-tests or Mann-Whitney tests (\**P* < 0.05; \*\**P* < 0.01; \*\*\**P* < 0.001). Distinct letters represent significant differences for one-way ANOVA or two-way ANOVA analyses.

## Data Availability

Mass-spectrometry-related data are deposited to the ProteomeXchange Consortium (https://proteomecentral.proteomexchange.org) via the iProX partner repository (Chen et al. 2022; Ma et al. 2019). IP–MS data: ProteomeXchange ID PXD059528; Proteomic sequencing data: Project ID IPX0013675000, ProteomeXchange ID PXD069244, accessible at https://www.iprox.cn/page/PSV023.html;?url=1760078369747UGpM with password LCyG. The data that support the findings of this study are also available in the supplementary material of this article.

## Acknowledgements

We thank Dr. Shiming Liu from Chinese Academy of Agricultural Sciences for providing Forrest seeds and Dr. Yongqing Jiao from Henan Agricultural University for providing PI88788 seeds. This work was supported by the National Key Research and Development Program of China (2023YFD1401000, 2023YFD1400400 to S.H.), National Natural Science Foundation of China (32488102 to X.F., 32301809 to D.W., and 32272478, 32102146 to S.H.), Science and Technology Development Plan Project of Jilin Province of China (20240602056RC to D.W.), Innovation Team Project of Northeast Institute of Geography and Agroecology, Chinese Academy of Sciences (2022CXTD03 to D.W.).

## Author Contributions

X.F. conceived and directed the project. S.Y. guided the details of the project and revised the manuscript. D.W. and S.H. designed and performed most of the experiments. D.W., S.H., S.Y. and X.F. drafted the manuscript with approval of the version to be submitted from all authors. L.W., L.W. and H.Y. performed wet-lab experiments. L.C., Q.L. and M.H. conducted nematode-inoculation experiments. C.L. provided SCNs. All authors read and approved of its content.

## Supporting Information legends

Supplemental Figure S1. Confocal-microscopy analysis of the physical interactions between SNAP18 isoforms and NSF.

Supplemental Figure S2. *SNAP18* expression remains stable before and after SCN infection.

Supplemental Figure S3. SNAP18 isoform abundance in transiently transgenic *N. benthamiana* leaves.

Supplemental Figure S4. The *lmm3* mutant is associated with increased autophagic activity rather than starvation-induced initiation.

Supplemental Figure S5. qRT–PCR of autophagy markers in 8-d-old Williams 82, *lmm3* and heterozygous *LMM3^+/-^* seedlings.

Supplemental Figure S6. Comparative proteomic analysis of *lmm3* and Williams 82 soybean roots under SCN-infected and uninfected conditions.

Supplemental Figure S7. NSF negatively impacts SCN resistance.

Supplemental Figure S8. Protein alignment of soybean-derived SNAP18 and NSF (Glyma.07G195900) with α-SNAP and NSF from *Rattus norvegicus* (PDB ID code 3J97, 20S complex crystal structure).

Supplemental Figure S9. Proposed model of SNAP18_lmm3_ autophagic degradation mediated by ATG8-driven autophagy.

Supplemental Figure S10. Amino-acid sequences of SCN-resistance-associated α-SNAP proteins.

Supplemental Table S1. Differential abundance analysis of core autophagy-related proteins in *lmm3* and Williams 82 roots under mock and SCN-infected conditions.

Supplemental Table S2. Partial list of representative proteins identified via IP–MS using SNAP18–GFP as bait.

Supplemental Table S3. PDBePISA analysis of the interaction structures (SNAP18–NSF and SNAP18–ATG8f) predicted by AlphaFold3.

Supplemental Table S4. Primers used in this study.

## Supporting Information

**FIGURE S1.**
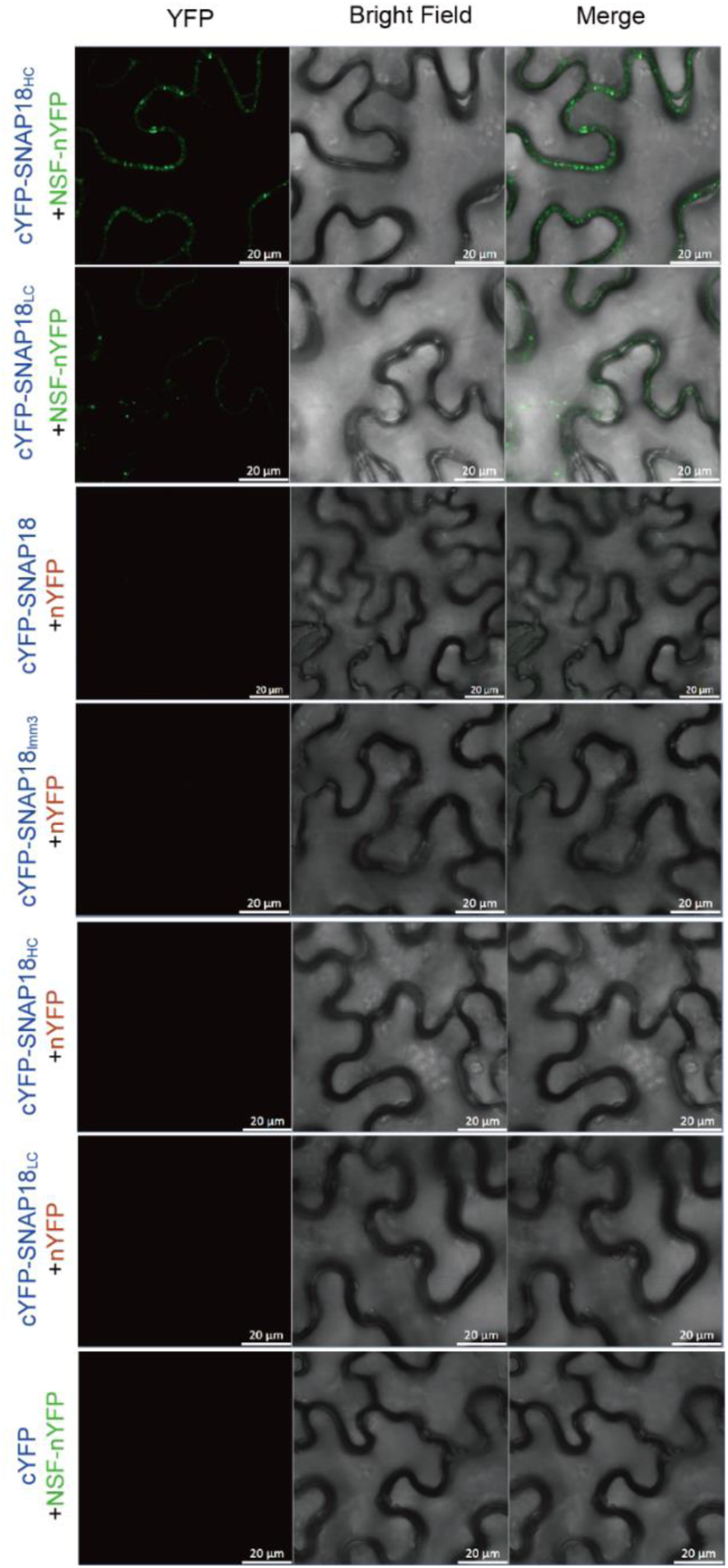
Confocal-microscopy analysis of the physical interactions between SNAP18 isoforms and NSF. BiFC assays in *N. benthamiana* demonstrate weakened binding between SNAP18HC and SNAP18LC with NSF. Negative-control co-expression experiments for Figure 1B are included. Three independent experiments were performed, yielding similar results. Scale bar = 20 μm.

**FIGURE S2.**
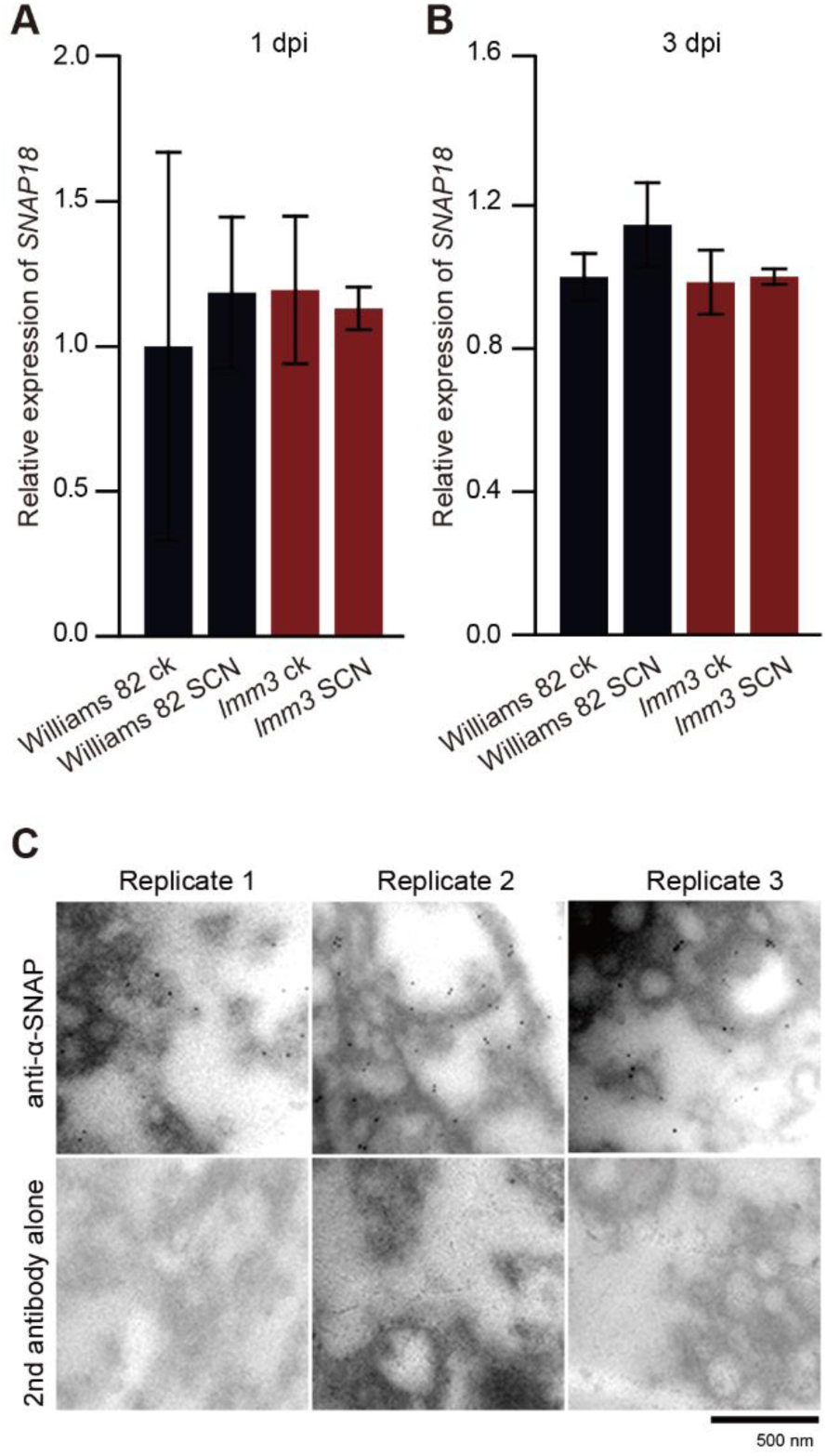
*SNAP18* expression remains stable before and after SCN infection. (A, B) *SNAP18* expression in roots of untreated control (‘ck’) Williams 82 and *lmm3* mutants at 1 dpi (A) or 3 dpi (B) following SCN infection (‘SCN’). Expression levels are normalized to the Williams 82 control and show the average from three biological replicates ±SD. Statistical analysis was performed by two-way ANOVA. (C) At 4 dpi, an electron micrograph (with brightness adjustment) reveals immunogold-labeled SNAP18lmm3 in syncytial cells of the *lmm3* mutant. The negative control consisted of the secondary antibody alone. Solid black particles indicate immunogold particles. Three biological replicates are presented. Scale bar = 500 nm.

**FIGURE S3.**
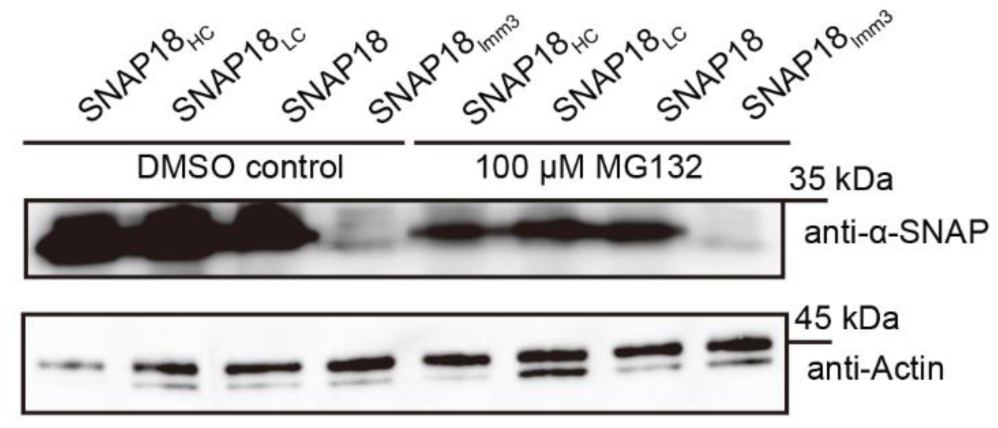
SNAP18 isoform abundance in transiently transgenic *N. benthamiana* leaves. Transient expression of untagged SNAP18, SNAP18HC, SNAP18LC and SNAP18lmm3 under the CaMV *35S* promoter. Leaves were treated with 100 μM MG132 or DMSO as a mock control. Protein levels were analyzed by western blot with actin serving as a loading control. The experiment was repeated three times with consistent results.

**FIGURE S4.**
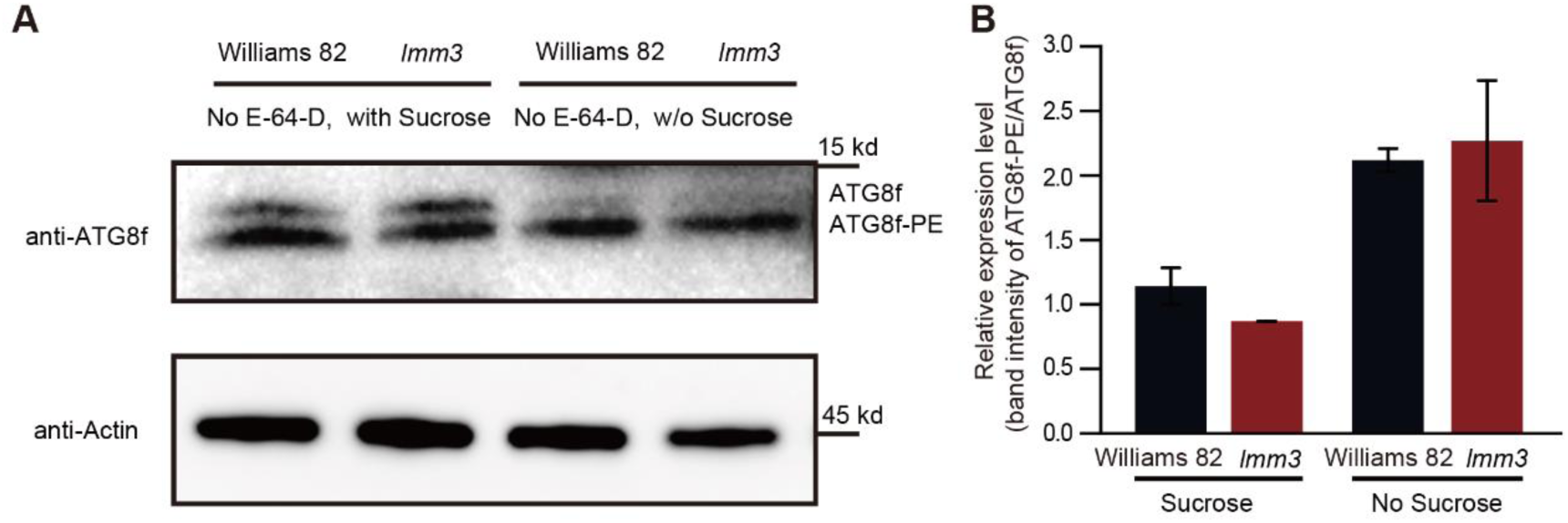
The *lmm3* mutant is associated with increased autophagic activity rather than starvation-induced initiation. (A) Western blot of ATG8f abundance in 7-d-old root extracts from Williams 82 and *lmm3* mutants. Seeds were germinated for 3 d before the roots were transferred to treatment conditions for an additional 4 d. Sucrose treatment consisted of Murashige and Skoog (MS) medium supplemented with 30 g/L sucrose, while sucrose starvation was performed by placing roots on MS medium without additional carbon sources. Samples were treated without 100 μM E-64-D. Actin was used as the loading control. (B) Ratio of ATG8f–PE to ATG8f based on (A). Data are the mean ±SEM of n = 2. Two-way ANOVA.

**FIGURE S5.**
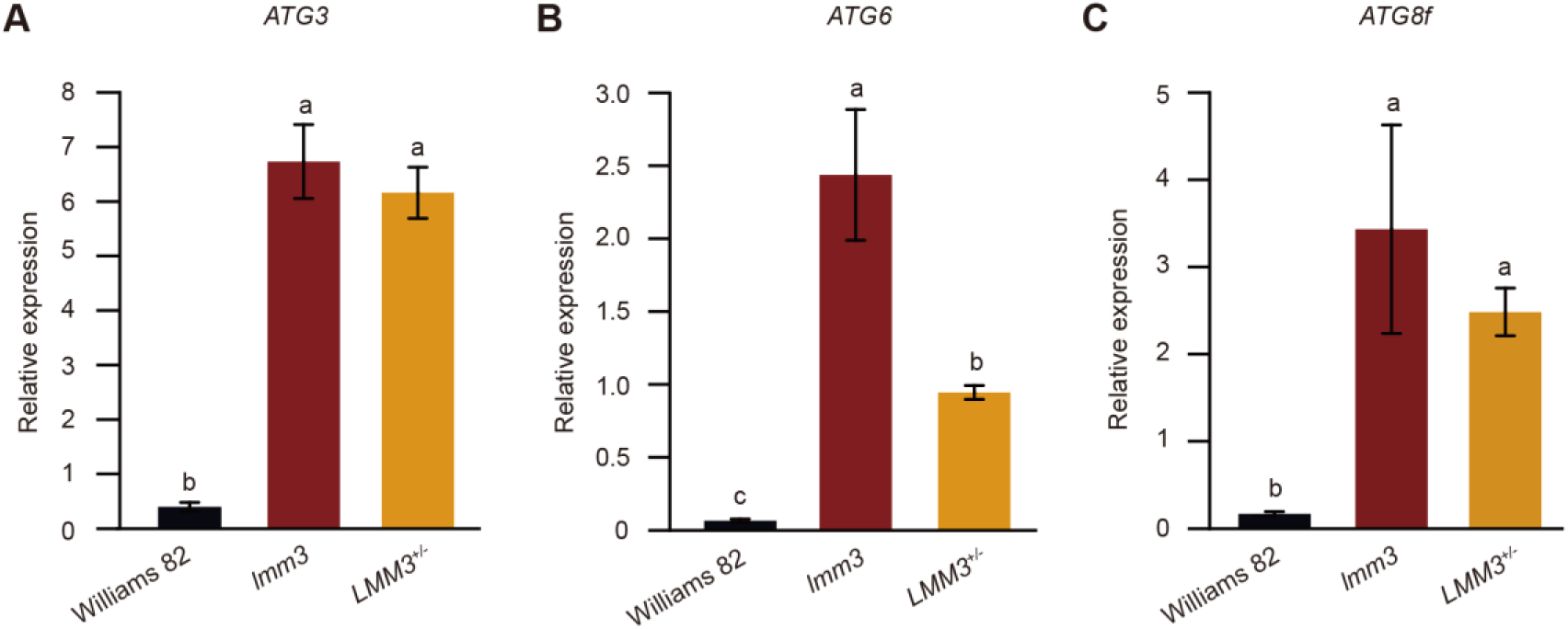
qRT–PCR of autophagy markers in 8-d-old Williams 82, *lmm3* and heterozygous *LMM3^+/-^*seedlings. Relative expression was normalized to *ACT11*. Data are as the mean ±SD of three biological replicates. One-way ANOVA, different letters represent *P* < 0.05.

**FIGURE S6.**
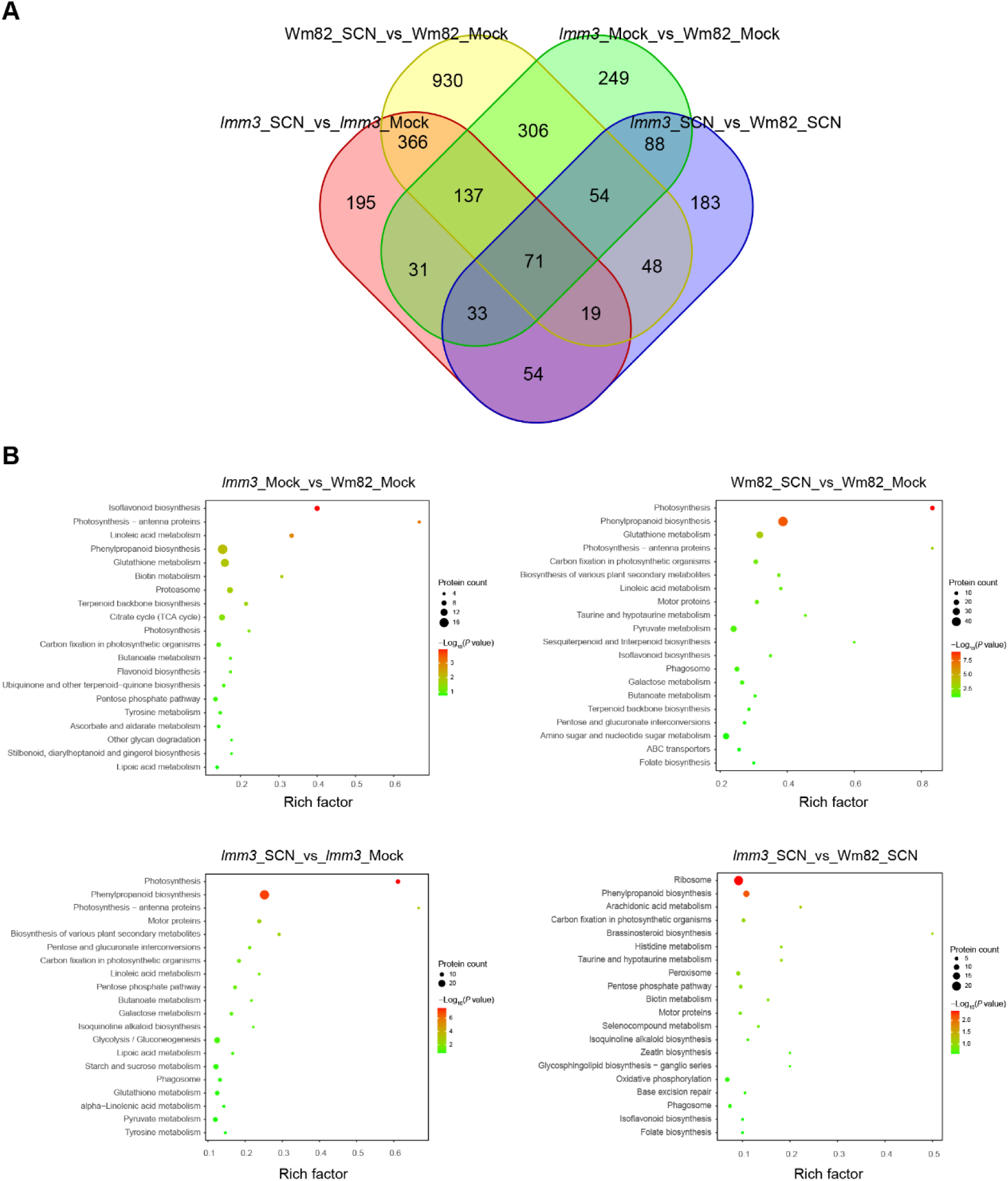
Comparative proteomic analysis of *lmm3* and Williams 82 soybean roots under SCN-infected and uninfected conditions. (A) Venn diagram showing the number of differentially expressed proteins (DEPs) across four comparison groups, integrating genotypic differences (*lmm3* vs. Williams 82) and SCN infection status (uninfected/mock vs. infected): (1) Wm82_SCN vs Wm82_Mock; (2) *lmm3*_Mock vs Wm82_Mock; (3) *lmm3*_SCN vs *lmm3*_Mock; (4) *lmm3*_SCN vs Wm82_SCN. Overlapping regions indicate shared DEPs. (B) KEGG pathway enrichment bubble plots for each comparison group. The x axis represents the Rich factor (ratio of DEPs annotated to a pathway relative to the total number of proteins in that pathway), the y axis lists enriched KEGG pathways, bubble size corresponds to the number of DEPs in the pathway, and color intensity indicates statistical significance (–log₁₀ *P*-value).

**FIGURE S7.**
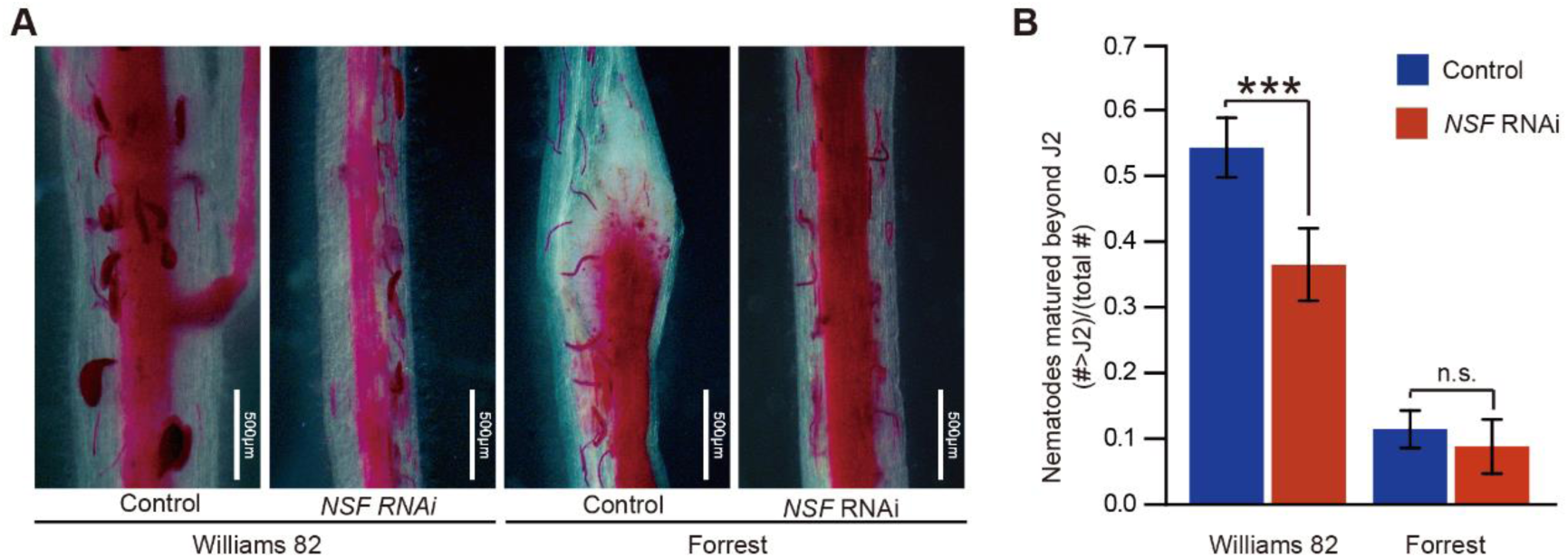
NSF negatively impacts SCN resistance. (A) Representative fuchsin-stained *H. glycines* at 12 dpi on transgenic hairy roots expressing either tdtomato (control) or *HIPpro:NSF* RNAi constructs with tdtomato as a selection marker in Williams 82 and Forrest backgrounds. Scale bar = 500 μm. (B) Analysis of SCN development beyond J2 stage in transgenic hairy roots expressing the transformation control or NSF RNAi constructs in the Williams 82 and Forrest backgrounds as in (A). Data are the mean ±SD of n ≥ 9 hairy roots. \*\*\**P* < 0.001, Student’s *t*-test.

**FIGURE S8.**
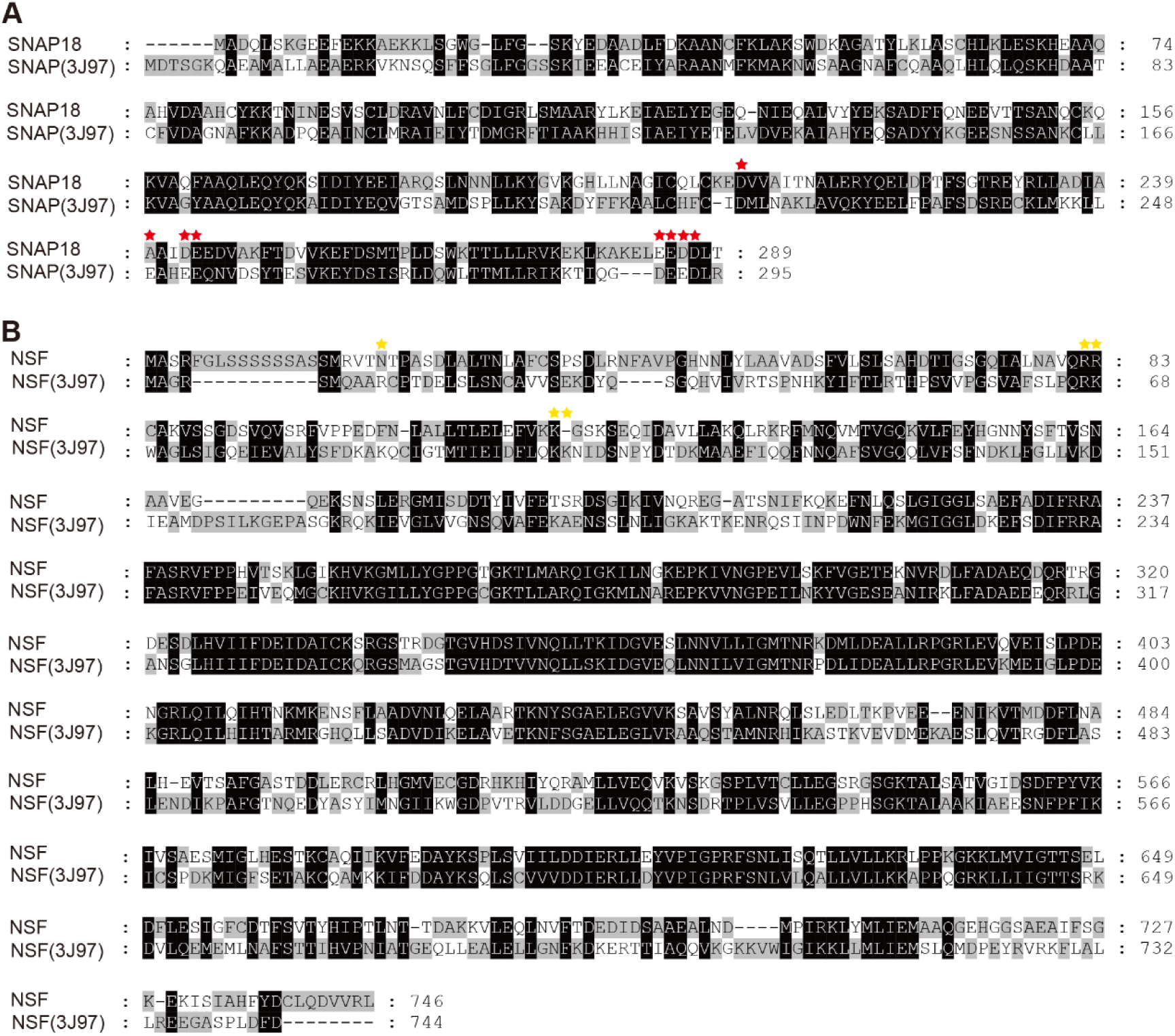
Protein alignment of soybean-derived SNAP18 and NSF (Glyma.07G195900) with α-SNAP and NSF from *Rattus norvegicus* (PDB ID code 3J97, 20S complex crystal structure). (A, B) Alignment of α-SNAP protein sequences (A) and alignment of NSF protein sequences (B). Red and yellow stars indicate key residues involved in α-SNAP and NSF binding within the crystal structure. Alignments were performed using the Clustal W algorithm in MEGA (version 5.05) and visualized with GeneDoc.

**FIGURE S9.**
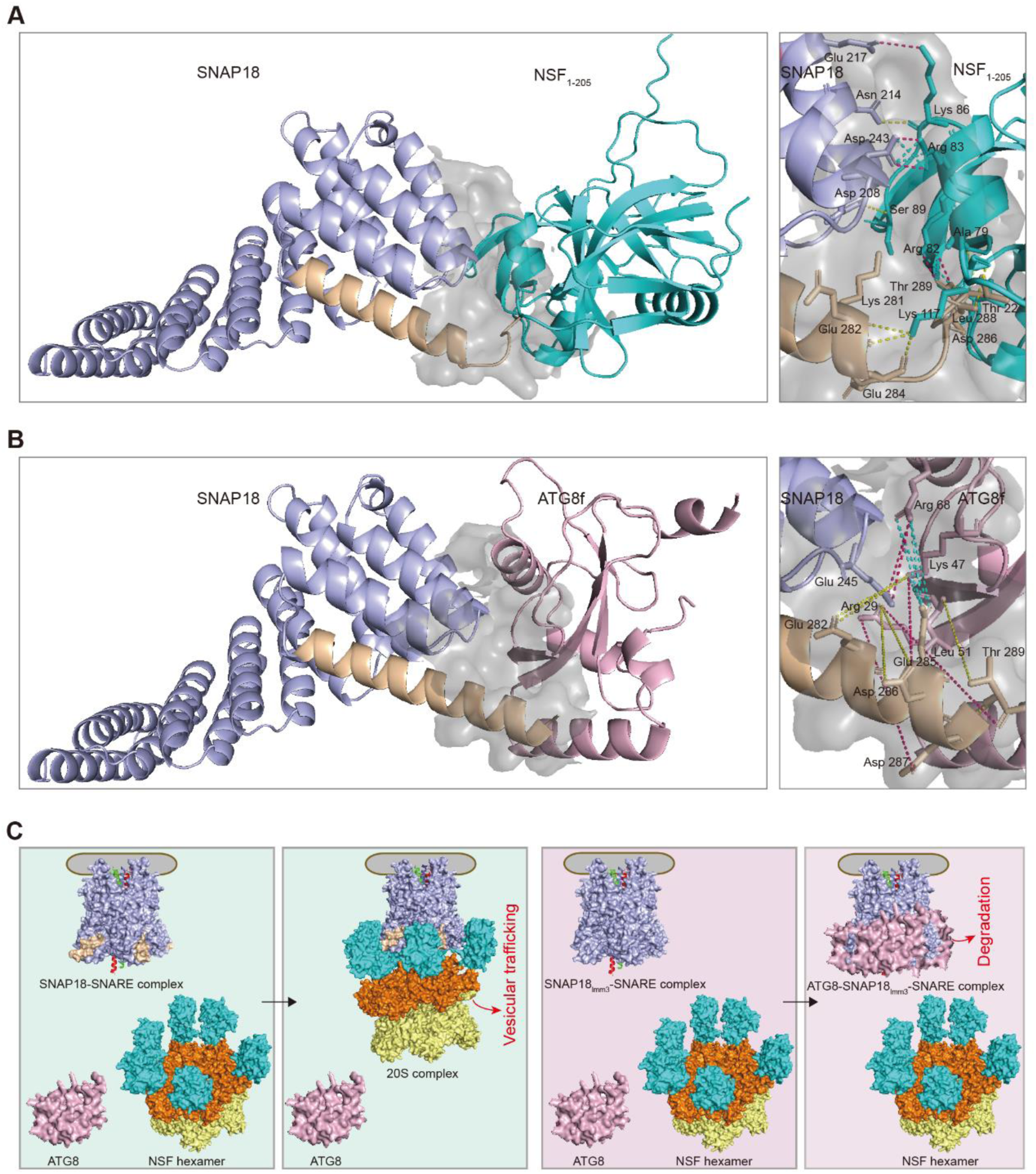
Proposed model of SNAP18lmm3 autophagic degradation mediated by ATG8-driven autophagy. (A, B) Predicted structures of SNAP18–NSF (A) and SNAP18–ATG8f (B) interactions determined by AlphaFold3. The predicted structures are displayed as cartoons. SNAP18 is shown in light blue, with the wheat-colored domain representing the C-terminal truncation of 24 amino acids encoded by *lmm3*. NSF is shown in cyan, with only the N-terminal 205 amino acids (NSF1-205) displayed. ATG8f is shown in light pink. Interaction interfaces between the two proteins are highlighted in light gray. Enlarged views of these interfaces are shown in the right panel, with residues involved in direct interactions displayed as sticks. Hydrogen bonds (yellow dotted lines), salt bridges (cyan dotted lines), and combined hydrogen bonds and salt bridges (hot pink dotted lines) are annotated. Interface interactions in the predicted structures were evaluated using PDBePISA (https://www.ebi.ac.uk/msd-srv/prot_int/pistart.html). (C) Proposed model for the autophagic-degradation mechanism for SNAP18lmm3. Left (light-green background): The NSF hexamer associates with the SNAP18-SNARE complex, resulting in the formation of a 20S complex. In this configuration, there is no available binding site for ATG8 on the complex. Consequently, SNAP18 remains unaffected by ATG8-dependent autophagy and is not degraded. Right panel (light-pink background): In the *lmm3* mutant, the SNAP18lmm3–SNARE complex is deficient in its C-terminal, which hinders its interaction with the NSF hexamer. This absence of the interaction with NSF enables ATG8 to readily associate with truncated SNAP18lmm3, facilitating its autophagic degradation via the ATG8-mediated autophagy pathway.

**FIGURE S10.**
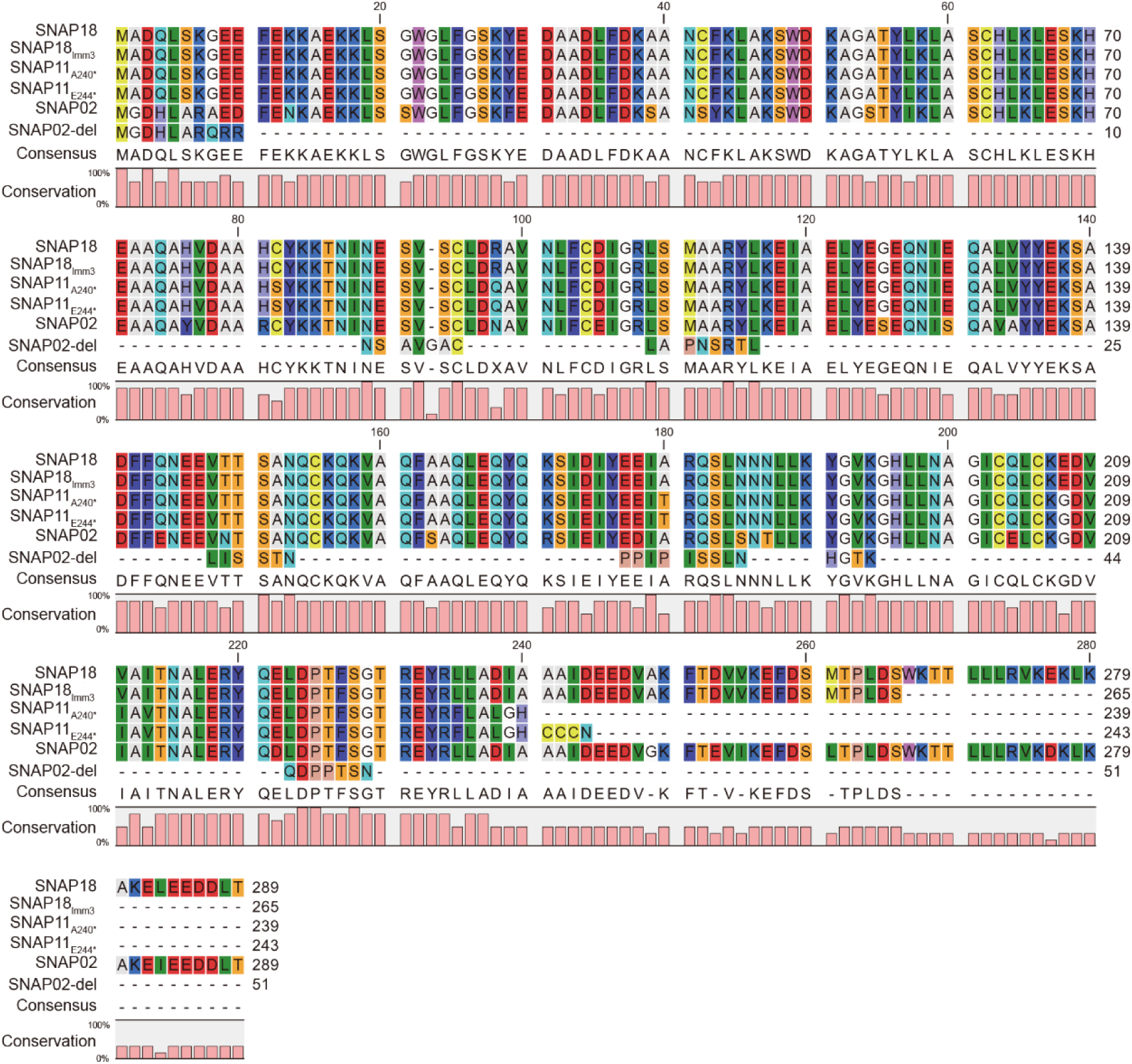
Amino-acid sequences of SCN-resistance-associated α-SNAP proteins. The alignment includes six known soybean α-SNAP variants: SNAP18 (289 aa) from Williams 82; SNAP18lmm3 (265 aa) from the *lmm3* mutant in the Williams 82 background; two truncated versions of SNAP11, namely SNAP11A240* (239 aa) and SNAP11E244* (243 aa) found in Forrest and Peking SCN-resistant-type accessions; SNAP02 (289 aa) from Williams 82; and SNAP02-del (51 aa), originating from PI 437654. Alignments were performed using the L-INS-i (accurate) option in Mafft and visualized with CLC Sequence Viewer 8.0.

**TABLE S1.** Differential abundance analysis of core autophagy-related proteins in *lmm3* and Williams 82 roots under mock and SCN-infected conditions.

| GeneID | Description | FC(*lmm3*_SCN_vs_<br>*s_lmm3*_Mock) | FC(Wm82_SCN_vs<br>_Wm82_Mock) | FC(*lmm3*_Mock_<br>vs_Wm82_Mock) | FC(*lmm3*_SCN_<br>vs_Wm82_SCN) |
| --- | --- | --- | --- | --- | --- |
| Glyma.02G133400 | ATG2 | 1.49 | 1.18 | 0.86 | 1.08 |
| Glyma.07G038100 | ATG12 | 1.45 | 0.94 | 1 | 1.53 |
| Glyma.09G231000 | ATG3 | 1.47 | 0.84 | 0.55 | 0.97 |
| Glyma.11G233000 | ATG27 | 1 | 1.46 | 1.45 | 0.99 |
| Glyma.12G005700 | ATG3 | 0.71 | 0.44 | 0.92 | 1.47 |
| Glyma.14G048000 | ATG16 | 1.38 | 2.43 | 1.53 | 0.87 |
| Glyma.16G019300 | ATG16 | 1.37 | 2.18 | 3.08 | 1.94 |

**TABLE S2.**
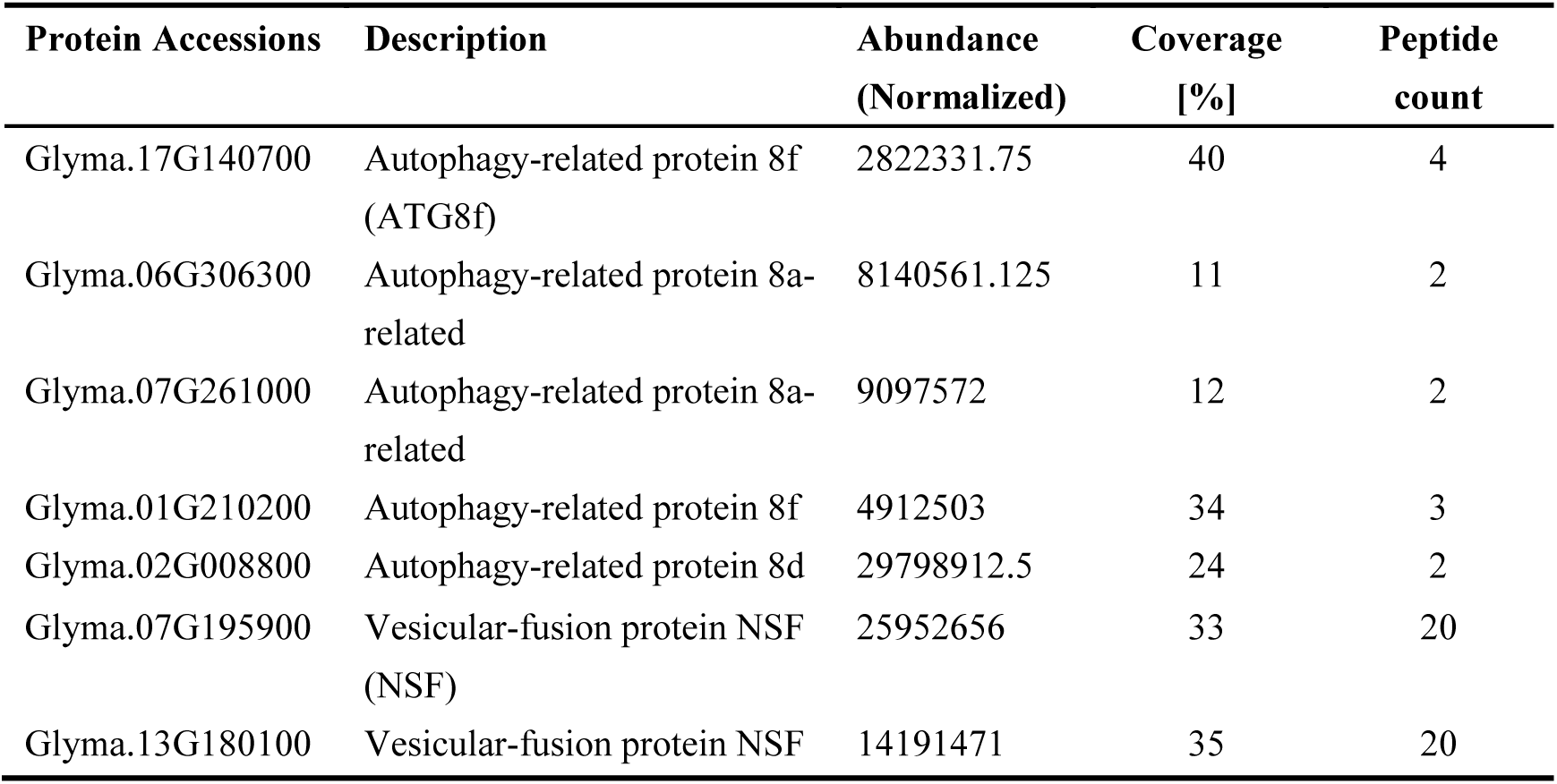
Partial list of representative proteins identified via IP–MS using SNAP18–GFP as bait.

**TABLE S3.**
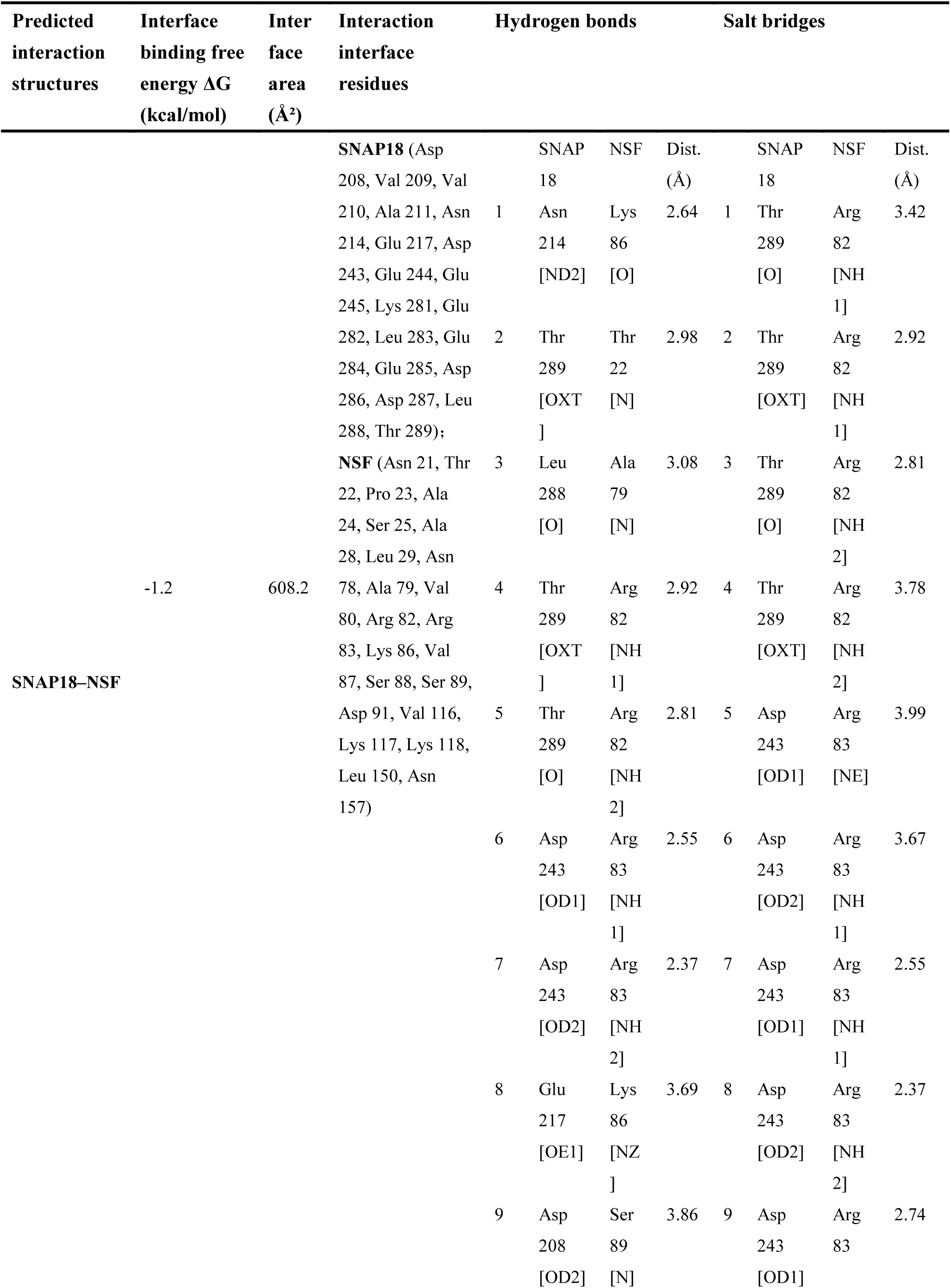
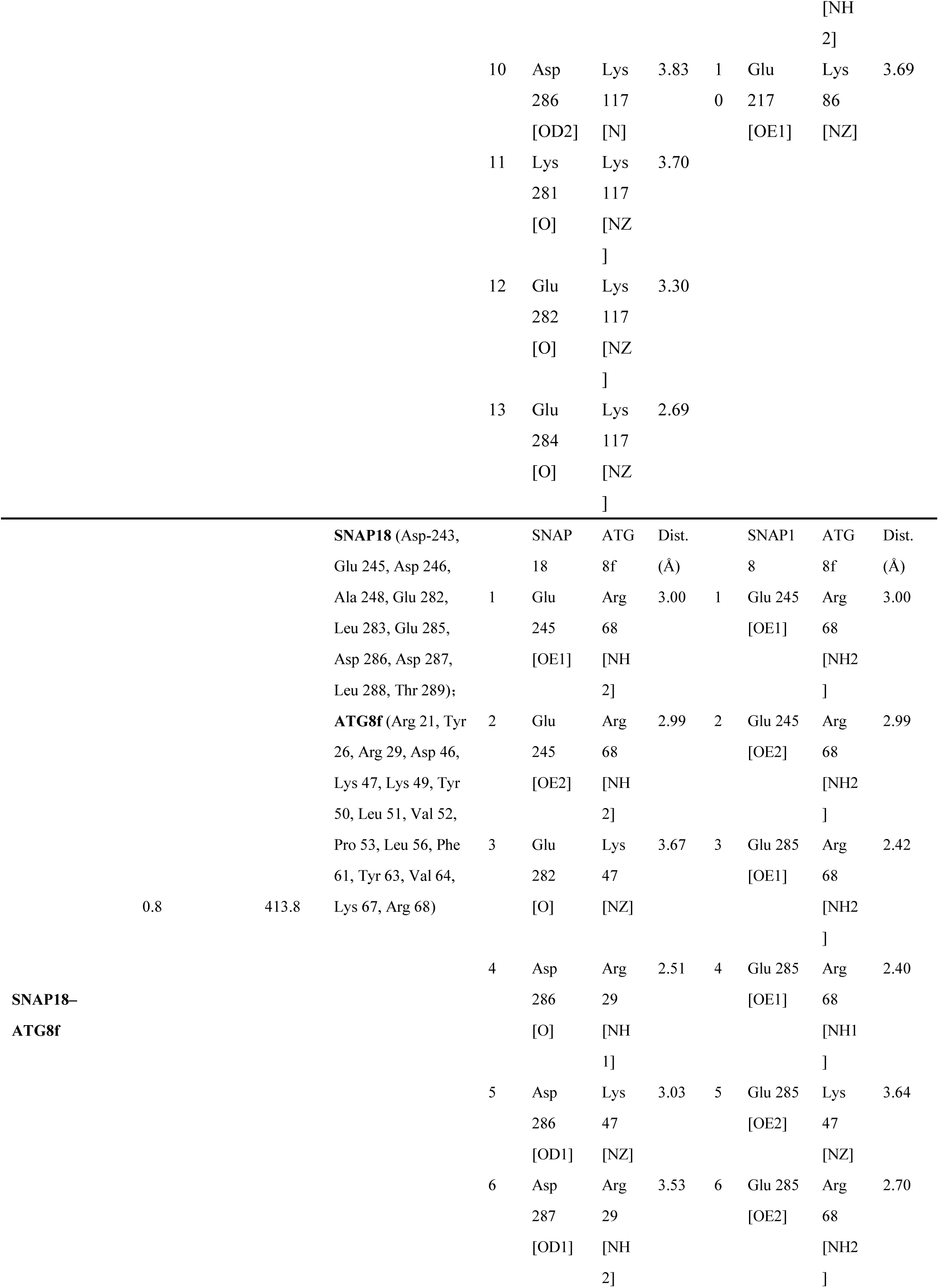
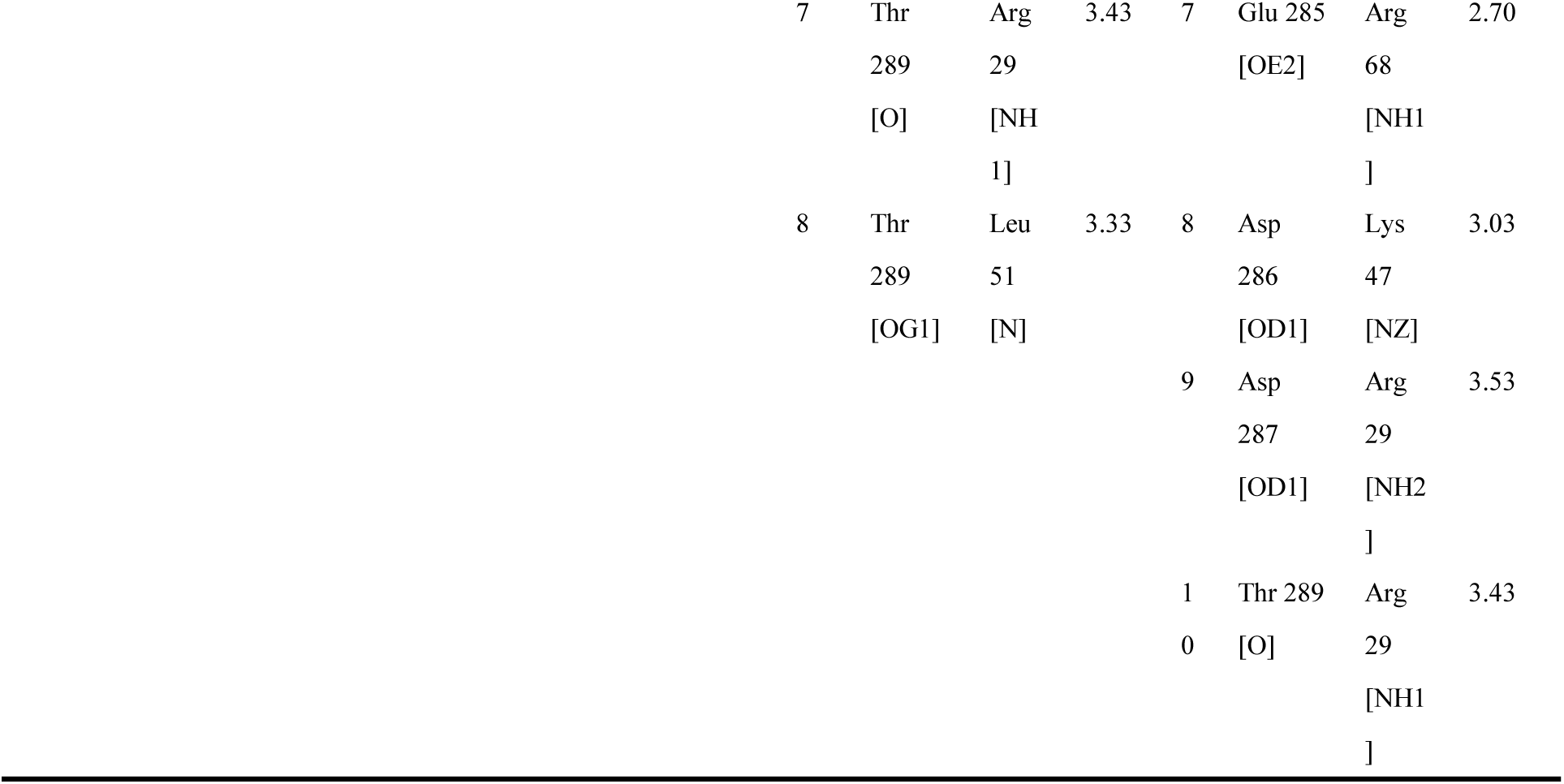
PDBePISA analysis of the interaction structures (SNAP18–NSF and SNAP18–ATG8f) predicted by AlphaFold3.

**TABLE S4.**
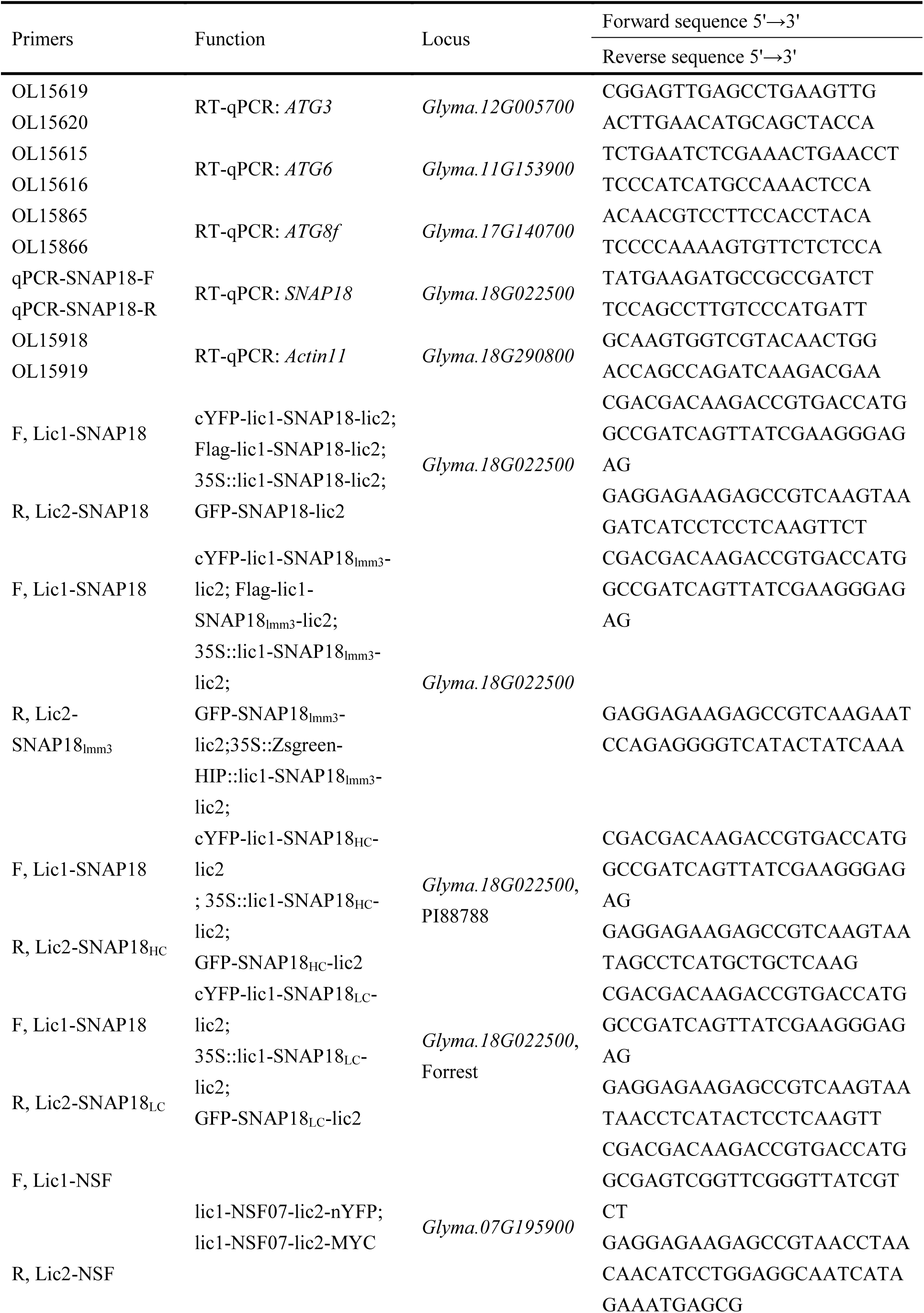
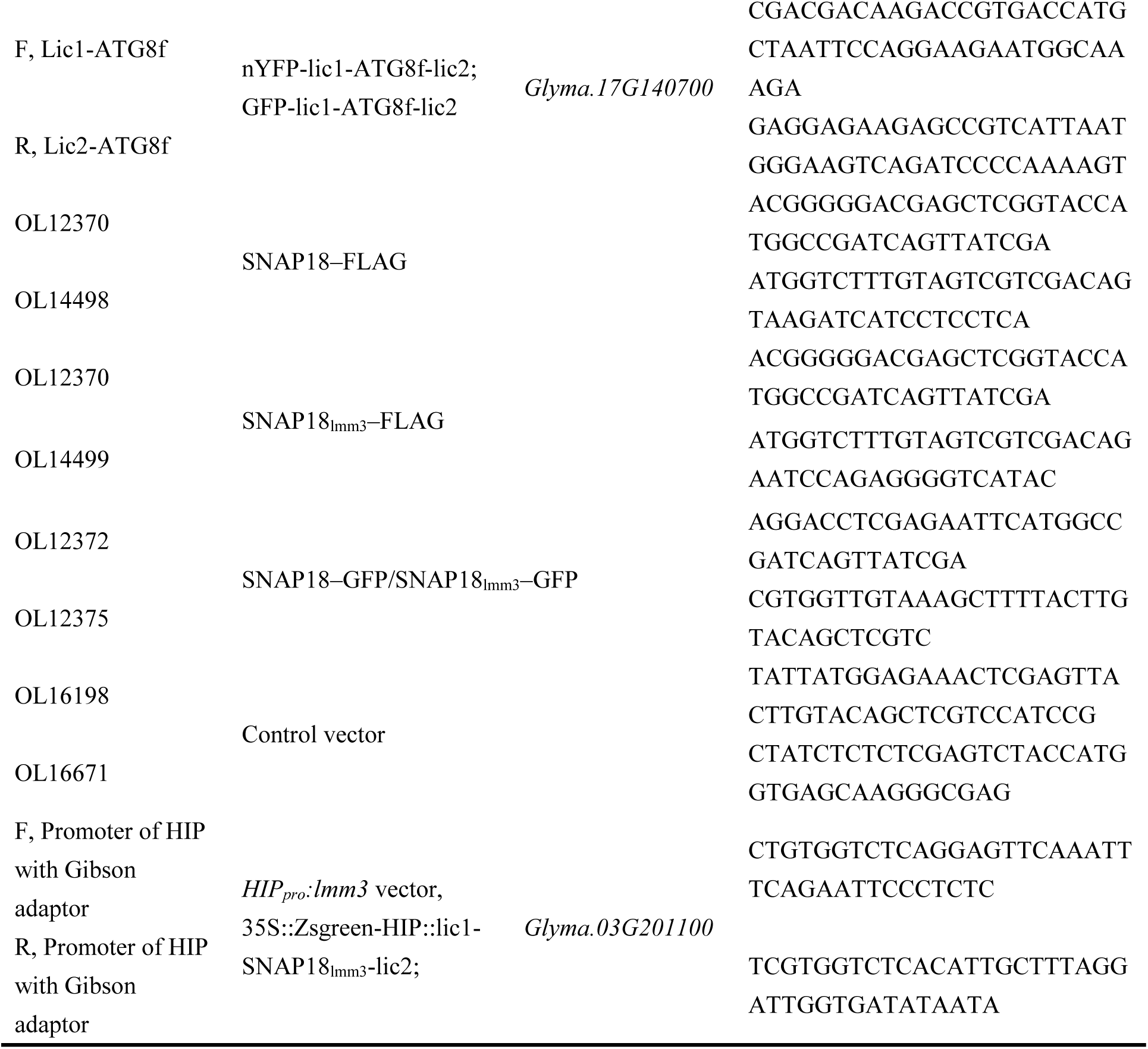
Primers used in this study.

